# Translational readthrough as a potential therapeutic for AIPL1-associated Leber Congenital Amaurosis in a patient-derived iPSC-retinal organoid model

**DOI:** 10.1101/2021.12.17.473147

**Authors:** Amy Leung, Almudena Sacristan-Reviriego, Pedro R. L. Perdigão, Hali Sai, Michalis Georgiou, Angelos Kalitzeos, Amanda-Jayne F. Carr, Peter J. Coffey, Michel Michaelides, James Bainbridge, Michael E. Cheetham, Jacqueline van der Spuy

## Abstract

Leber Congenital Amaurosis type 4 (LCA4), caused by *AIPL1* mutations, is characterised by severe sight impairment in infancy and rapidly progressive degeneration of photoreceptor cells. We generated retinal organoids using induced pluripotent stem cells (iPSCs) from renal epithelial cells obtained from four children with *AIPL1* nonsense mutations. iPSC-derived photoreceptors exhibited the molecular hallmarks of LCA4, including undetectable AIPL1 and rod cGMP phosphodiesterase (PDE6) compared to control or CRISPR corrected organoids. Moreover, increased levels of cGMP were detected. The translational readthrough inducing drug (TRID) PTC124 was investigated as a potential therapeutic. LCA4 retinal organoids exhibited rescue of AIPL1 and PDE6; however, the level of full-length, functional AIPL1 induced through PTC124 treatment was insufficient to reduce cGMP levels and fully rescue the LCA4 phenotype. LCA4 retinal organoids are a valuable platform for the *in vitro* investigation of the molecular mechanisms that drive photoreceptor loss and for the evaluation of novel therapeutics.

## Introduction

Leber Congenital Amaurosis (LCA), the most severe form of inherited retinal degeneration (IRD), is characterized by the early and progressive severe loss of vision within the first few years of life^1^. LCA patients typically present with nystagmus, amaurotic pupils and markedly reduced/absent full-field electroretinograms (ERG).^2^ LCA is genetically heterogeneous, with 26 genes associated with the disease (Retinal Information Network https://sph.uth.edu/retnet), and is typically inherited in an autosomal recessive manner. Biallelic mutations in the aryl hydrocarbon receptor interacting protein-like 1 *(AIPL1)* gene (LCA type 4; OMIM 604392) are associated with clinical features that are at the severe end of the spectrum of LCA and account for up to 5-10% of LCA cases.^3–5^

The 384 amino acid protein AIPL1, exclusively expressed in retinal photoreceptors and pineal gland,^3,6,7^ has a FK506-binding protein (FKBP)-like domain at the N terminus, followed by a tetratricopeptide repeat (TPR) domain and a C-terminal primate-specific proline rich domain (PRD). AIPL1 acts as a specialized molecular co-chaperone, together with HSP90, enabling the correct folding and assembly of the guanosine-3’,5’-cyclic monophosphate (cGMP)-specific phosphodiesterase 6 (PDE6), a critical holoenzyme in the phototransduction cascade that hydrolyses cGMP in rods and cones upon light stimulation.^8,9^ It has been shown that the AIPL1 FKBP-like domain binds the isoprenylated PDE6 catalytic subunits,^10–12^ while the AIPL1 TPR domain interacts with the regulatory PDE6g subunit^13^ and the EEVD motif located at the C-terminus of HSP90.^14,15^ Several studies in mouse retina revealed that, with the reduction or absence of AIPL1, cone and rod PDE6 levels decrease^16–18^ and rod PDE6 subunits are misassembled and targeted to proteasomes for degradation.^10^ As a result of reduced PDE6 levels, cGMP accumulates leading to rapid photoreceptor degeneration in the knockout mouse model.^18^ Similarly, LCA-associated mutations compromising the integrity of the AIPL1 FKBP-like or TPR domain failed to efficiently modulate rod or cone PDE6 function *in vitro*.^9,19,20^

Currently, there is no cure or treatment for AIPL1-associated LCA (LCA4), which has a devastating impact on the quality of life. To date, replacement gene therapy is the most promising potential therapy for LCA. Landmark clinical trials involving adult patients with *RPE65* reported some improvement of visual function and no serious complications.^21,22^ This led to the approval of the first ocular gene therapy, voretigene neparvovec-rzyl (Luxturna), for the treatment of RPE65-associated LCA. The potential of gene therapy to treat LCA caused by *AIPL1* mutations has been examined in preclinical studies conducted in mouse models of AIPL1 deficiency. These studies reported a delayed retinal degeneration and rescue of photoreceptor function following delivery of the *AIPL1* gene through adeno-associated virus (AAV).^23–25^ Several novel stem cell-based therapies are currently under development.^26^ The transplantation of cones derived from mouse embryonic stem cells (mESCs) or human pluripotent stem cells (hPSCs) into the *Aipl1-1*-mouse model resulted in the regeneration of the photoreceptor layer and synaptic-like structures post-transplantation,^27,28^ suggesting the potential of the stem-cell approach to restore vision in *AIPL1*-LCA patients.

Nonsense variations giving rise to premature termination codons (PTCs) are extensively found in genetically transmitted disorders, including *AIPL1-associated* LCA.^29^ Almost two thirds of all disease-associated variations in *AIPL1* encode a nonsense variant. The *in vitro* expression of *AIPL1* c.94C>T (p.R32X), c.216G>A (p.W72X), c.264G>A (p.W88X), c.487C>T (p.Q163X), c.582C>G (p.Y194X), c.665G>A; (p.W222X) and c.834G>A (p.W278X) resulted in non-functional truncated protein products, thus confirming their disease-causing status.^15,20^ Translational readthrough inducing drugs (TRIDs) promote ribosomal misreading of PTCs and restore the production of full-length functional proteins.^30–33^ Among all the TRIDs, PTC124 (also known as Ataluren or Translarna™), is the only one that has been authorized for clinical use in Duchenne muscular dystrophy (DMD) following clinical trials.^34^ PTC124, with minimal ocular side effects, has also been investigated as a potential TRID in several hereditary ocular disorders including Usher syndrome,^35,36^ choroideremia,^37,38^ retinitis pigmentosa,^39–41^ LCA^42^ and congenital aniridia.^43–45^ These findings suggest that TRIDs could rescue nonsense *AIPL1* mutations causing LCA.

Reprogramming patient-derived somatic cells into induced pluripotent stem cells (iPSCs) has revolutionized the study of genetic diseases in the last decade.^46^ Likewise, the investigation of human eye development has benefitted from advances in stem cell differentiation towards three dimensional (3D) retinal organoids (ROs). These 3D ROs contain all major retinal cells types and mature photoreceptors develop outer segment-like structures, connecting cilia, inner segments rich in mitochondria and synaptic connections.^47^ This technology provides the opportunity for modelling different IRDs *in vitro* and testing personalised treatments. In this study, we have developed and characterised the first model of *AIPL1*-LCA patient-derived ROs from renal epithelial cells carrying nonsense variations in the *AIPL1* gene.

We report that the patient-derived ROs display the key molecular hallmarks of LCA4, with absence of detectable AIPL1 and rod cGMP PDE6 subunits, and elevated cGMP levels in photoreceptor cells. Using this platform, we examined the efficacy of PTC124 induced translational readthrough in rescuing full-length AIPL1 in ROs harbouring the nonsense mutation c.834G>A, p.W278X. PTC124 was able to rescue full-length AIPL1 in rod and cone photoreceptor cells. Moreover, the rescue of rod PDE6 was observed in a small number of photoreceptors in ROs homozygous for c.834G>A p.W278X. However, the level of rescue was insufficient to reduce overall cGMP levels.

## Results

### Generation and characterisation of iPSCs from LCA4 patients harbouring different mutations in *AIPL1*

In order to generate iPSCs, renal epithelial cells were purified from urine samples from 4 different LCA4 patients aged 2-4 years (LCA4-1, LCA4-2, LCA4-3, LCA4-4) (Figure 1). The collection of urine is a minimally invasive procedure advantageous for obtaining somatic cells from young patients with paediatric disease. From this group of 4 patients, there are 3 unique mutation combinations at the *AIPL1* locus (Figure 1A), with all patients carrying at least one copy of the c.834G>A, p.W278X mutation. Patient LCA4-1 is homozygous for c.834G>A, p.W278X. The patient had nystagmus from birth and poor visual function. On examination at 3.2 years old, the patient had a best corrected visual acuity (BCVA) of perception of light in both eyes with bilateral roving eye movements and horizontal and vertical nystagmus. Optical coherence tomography (OCT) revealed residual foveal outer retinal structure (Supplementary Figure 1A). Patients LCA4-2 and LCA4-3 are compound heterozygous for c.834G>A, p.W278X and a mutation in intron 3 that abolishes the putative intronic splice acceptor site, c.466-1G>C. The patients are monozygotic twins, born from non-consanguineous parents, and were symptomatic at birth, exhibiting poor visual behaviour and nystagmus. Fundi were abnormal with generalised reduction in arterial calibre and macular pigmentary disturbance, with normal retinal periphery. Electroretinograms performed at the age of 10 months showed extinguished photopic and scotopic responses in both infants. BCVA was light perception for both patients and pupillary light reflex was present, but of weak amplitude. OCT performed at 2 years of age also revealed residual foveal outer retinal structure (Supplementary Figure 1B-C). Patient LCA4-4 is compound heterozygous for c.834G>A, p.W278X and c.665G>A, p.W222X. The clinical findings for LCA4-4 at 3 years and follow-up at 5 years of age have been previously described.^20^ The locations of these mutation sites in the *AIPL1* gene are highlighted in Figure 1B.

**Figure 1:**
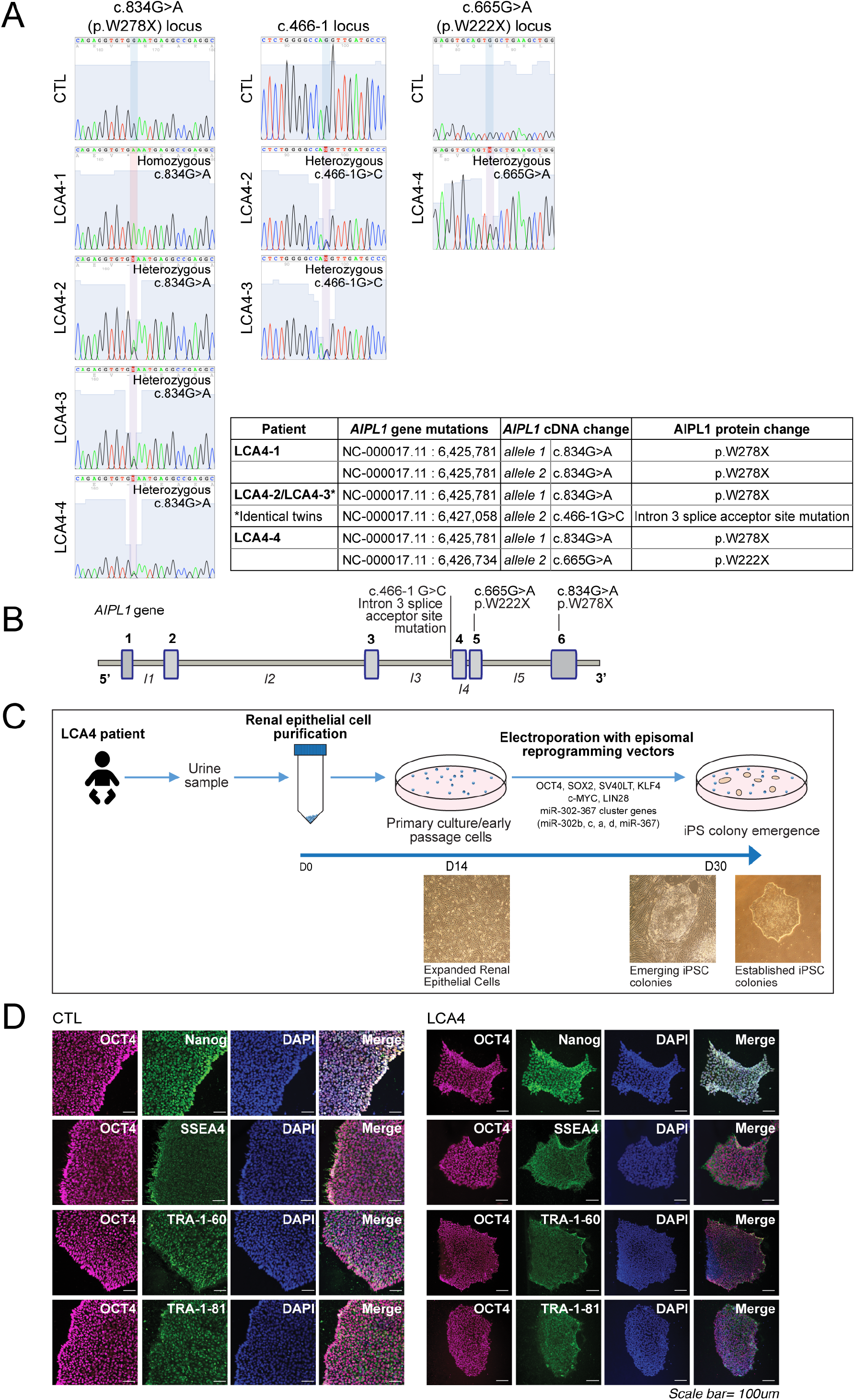
Characterisation of LCA4 patient mutations and iPSC generation from patient renal epithelial cells. A) Sequence analysis of the *AIPL1* gene in LCA4 patient samples. Sequence chromatograms of control (CTL) and LCA4 patient samples at the c.834G>A (p.W278X), c.466-1G>C and c.665G>A (p.W222X) loci. The table summarises the AIPL1 mutations (genomic site, transcript change, predicted protein change) present in LCA4 patients. B) Schematic diagram of *AIPL1* gene; the LCA4 patient mutation sites are highlighted. C) Schematic diagram and brightfield images detailing the timeline of iPSC generation from patient renal epithelial cells. Representative brightfield images of renal epithelial cells, emerging and established iPSC cultures from LCA4-1, LCA4-2, LCA4-3 and LCA4-4 are shown in Supplementary Figure 2A. D) IF analysis of CTL and LCA4-2 iPSCs for pluripotency markers (OCT4, Nanog, SSEA4, TRA-1-60, TRA-1-81). IF images for LCA4-1, LCA4-3 and LCA4-4 iPSC are shown in Supplementary Figure 2B. DAPI staining is in blue. Scale bars = 100μm.

iPSC generation was carried out as previously described (Figure 1C).^48^ In brief, primary renal epithelial cells were expanded for 14 days and then electroporated with episomal reprogramming vectors. Emerging iPSC colonies were visible approximately 2 weeks later, and these were selectively isolated and expanded to form multiple iPSC lines from all LCA4 patient cultures (Supplementary Figure 2A). Multiple iPSC lines from well-characterised controls (CTL)^49^ were expanded in parallel. Immunofluorescence (IF) labelling of iPSC cultures from CTL and LCA4 lines was carried out to confirm that the cells uniformly express pluripotency markers such as OCT4, NANOG, TRA1-80, TRA1-61 and SSEA4, as expected (Figure 1D, Supplementary Figure 2B). Gene expression analyses of early iPSC differentiation cultures towards ectodermal, mesodermal and endodermal lineages confirmed expression of the appropriate lineage markers, demonstrating trilineage differentiation potential (Supplementary Figure 2C).

### Characterisation of retinal organoid structure and retinal cell populations from LCA4-iPSCs

Retinal organoids (ROs) were derived from iPSCs as previously described.^28^ Retinoic acid (RA) was removed from the culture medium from day 100 onwards to further the photoreceptor (PR) maturation process (Figure 2A). Clearly defined, neuro-retinal vesicle (NRV) structures emerged after 3-4 weeks of culture from the monolayer differentiation, and displayed laminated photoreflective properties. Over time, the mechanically isolated NRVs expanded in size, while maintaining a clearly laminated structure (the outer nuclear layer (ONL)) on the surface that is comprised of maturing photoreceptor (PR) cells. After 24-26 weeks of culture following NRV isolation, the ONL PR cells formed clear ciliary extensions and distal structures that form a dense brush border of presumptive PR inner and outer segment structures (Figure 2B, Supplementary Figure 3A).

**Figure 2:**
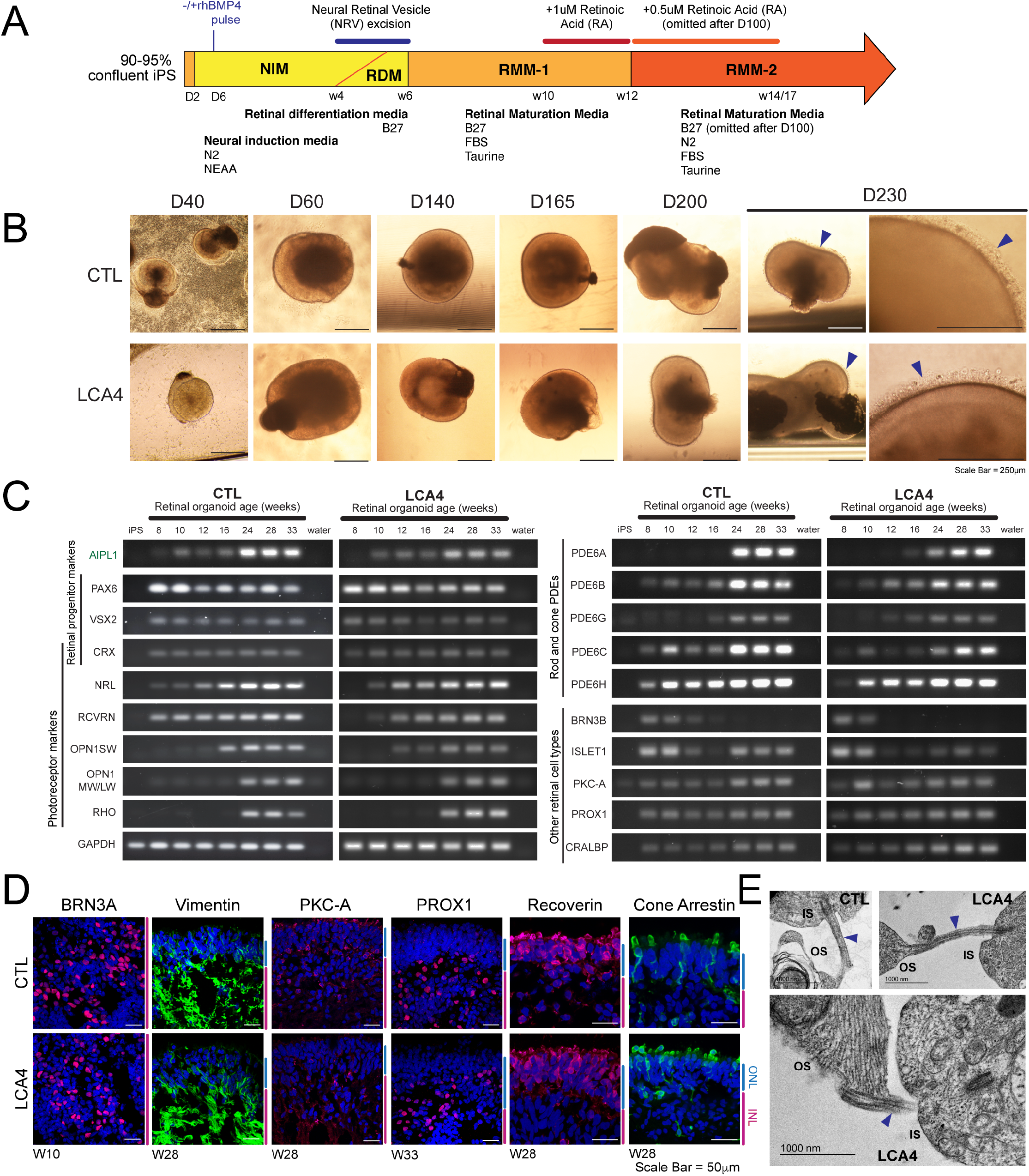
LCA4 iPSCs were able to generate retinal organoids (ROs) with all retinal cell types. A) Schematic diagram detailing the RO differentiation process. B) Brightfield images of developing CTL and LCA4-1 ROs (day (D) 40-230). A well-developed brush border of presumptive OS/IS photoreceptor structures was present on the surface of ROs by D230 (blue arrowheads). Brightfield images of RO differentiation for LCA4-2, LCA4-3 and LCA4-4 are shown in Supplementary Figure 3A. Scale bars = 250μm. C) Semi-quantitative PCR analysis of developing CTL and LCA4 ROs (W8-W33) for retinal development and retinal lineage markers. PCRs from LCA4-2 RO samples shown; the pattern of LCA4 RO gene expression was similar in all LCA4 patient lines. D) IF images of CTL and LCA4 ROs for retinal cell markers. BRN3A was expressed in retinal ganglion cells in early (W10) ROs. Vimentin (Muller glia), PKC-A (bipolar cells), PROX1 (amacrine/horizontal cells), Recoverin (PRs) and Cone arrestin (cones) were similarly distributed in CTL and LCA4 ROs. LCA4 images are representative of LCA4 ROs from all LCA4 patient lines. DAPI staining is in blue. ONL and INL regions highlighted at the side of images in blue and magenta. Scale bars = 50μm. E) Transmission electron microscopy images of CTL and LCA4 RO PR regions. Images from LCA4-1 are shown. Connecting cilia (blue arrowheads) were visible in both types of PRs. Outer segment (OS) structures contain membranous folds, and mitochondria were visible in inner segments (IS).

Gene expression analysis of W8-W33 CTL and LCA4 ROs for genes involved in early retinal development (*PAX6*, *VSX2*), photoreceptor development (*CRX*, *NRL*) and function (recoverin (*RCVRN*), rhodopsin (*RHO*), cone opsins (*OPN1SW*, *OPN1MW/LW*), phosphodiesterase subunits (*PDE6A*, *PDE6B*, *PDE6G*, *PDE6C*, *PDE6H)*) indicated that these genes were regulated in a similar temporal pattern in both CTL and LCA4 ROs (Figure 2C). The onset of expression of retinal progenitor markers and transcription factors was detected early on and sustained over time, with a noticeable increase in *NRL* expression with time. The expression of PR markers increased over time in both CTL and LCA4 RO, however a later onset of expression of rod and cone phototransduction components (*OPN1SW*, *OPN1MW/LW*, *RHO*, *PDE6A*, *PDE6B*, *PDE6G*)was observed. Interestingly, *AIPL1* expression was detectable in LCA4 ROs in a similar temporal pattern as CTL ROs with expression first detected at W8-W10 and with a noticeable increase at W24-W33, corresponding with the increased expression of retinal PR markers (including rod and cone *PDE6*). Similar patterns in CTL and LCA4 RO were also observed for genes associated with non-PR retinal cell types (Figure 2C). The expression of the retinal ganglion cell (RGC) marker *BRN3B* declined rapidly after expression early on, whereas RGC *ISL1* expression was biphasic with a dip in expression at W12-W16. The expression of markers for Muller glia (*CRALBP*), amacrine and horizontal (*PROX1*) and bipolar cells (*PKCA*) were sustained throughout in CTL and LCA4 ROs.

IF analysis of CTL and LCA4 RO sections confirmed that there was a similar pattern of expression and distribution of retinal cell markers within the ROs at varying stages of development (Figure 2D). BRN3A, which is expressed in RGCs (a cell population in early-staged ROs that gradually disappears as ROs develop), was abundant throughout the core of early-staged W10 ROs. In later-staged, mature ROs (W28-W33), which display a clearly delineated ONL-inner nuclear layer (INL) tissue structure, cells positive for vimentin (present throughout the Muller glia cell structure that extends into the ONL), PKCA (located in the cytoplasm of bipolar cells, spread throughout the INL), and PROX1 (transcription factor localised to the nuclei of horizontal and amacrine cells, located in the INL) are present in both CTL and LCA4 organoids. Examination of the PR markers RCVRN (expressed in both rod and cone populations) and Cone Arrestin (CARR; cones only) revealed strong expression of these markers in the cells within the ONL of both CTL and LCA4 ROs. Both RCVRN and CARR are distributed throughout the entirety of the PR cells, including the apical inner segment (IS) regions of the PRs. Examination of ROs for TUNEL positivity revealed that very few cells were TUNEL+ (any positive cells were mostly localised to the core of the RO and not the ONL PR), and patient RO do not appear to display changes to baseline levels of apoptosis (Supplementary Figure 3B). Transmission Electron Microscopy (TEM) also highlighted that the photoreceptors in LCA4 ROs are structurally similar to those see in CTL ROs, with clearly defined connecting cilia linking presumptive inner and outer segment structures of the PRs (Figure 2E). Overall, these data indicate that development of the RO retinal tissue structure and retinal cell types is comparable in CTL and patient ROs and lacking in overt indictors of neurodegeneration.

### All LCA4 ROs lack detectable AIPL1 protein, despite the presence of LCA4 *AIPL1* mRNA transcripts

ROs from all patient lines express *AIPL1* mRNA, as shown by PCR amplification of *AIPL1* (exon1-2 region) from W28 RO cDNA samples (Figure 3A). In order to determine the effect of the LCA4 mutations on *AIPL1* mRNA transcripts in ROs, the regions of interest (exons 3-5 for c.466-1G>C and c.665G>A, p.W222X; and exons 5-6 for c.834G>A, p.W278X) were amplified from W28 cDNA and sequenced (Figure 3B). The p.W222X transcript was not detectable in LCA4-4 RO cDNA, suggesting that the mRNA species undergoes nonsense mediated decay (NMD). NMD is generally efficient at degrading mRNA species with PTCs 50-55 base pairs upstream of exon-exon junctions, and c.665G>A, p.W222X is located 119 bp from the exon 5-6 junction. Sequencing of LCA4-2 and LCA4-3 samples revealed that the c.466-1G>C mutation, which ablates the intron 3 splice acceptor site, predominantly results in a transcript that is missing the first 24 base pairs of exon 4 (an in-frame 8 amino acid deletion; p.V156_Q163del). For all of the LCA4 RO samples the p.W278X transcript was detectable, indicating that this *AIPL1* mRNA species does not undergo or partially escapes NMD. This was expected as the mutation is located in the final *AIPL1* exon (exon 6), and PTCs in the final exons of genes often do not undergo substantial levels of NMD.^50^ The effects of the AIPL1 mutations in the 4 LCA4 ROs are summarised in Figure 3C, and the mutation sites in relation to the AIPL1 protein structure are highlighted in Figure 3D. The c.834G>A, p.W278X nonsense mutation codes for a premature stop codon expected to induce C-terminal truncation of the AIPL1 TPR domain, which is critical for the interaction with HSP90.^14,15^ Similarly, the c.665G>A, p.W222X mutation codes for a premature stop codon affecting the TPR domain, but the p.W222X transcript undergoes NMD resulting in the absence of translation of this protein species in patient ROs. The in-frame p.V156_Q163del mutation leads to the deletion of the linker between the FKBP-like and TPR domains, critical for the organisation and function of these domains relative to one another.^15^

**Figure 3:**
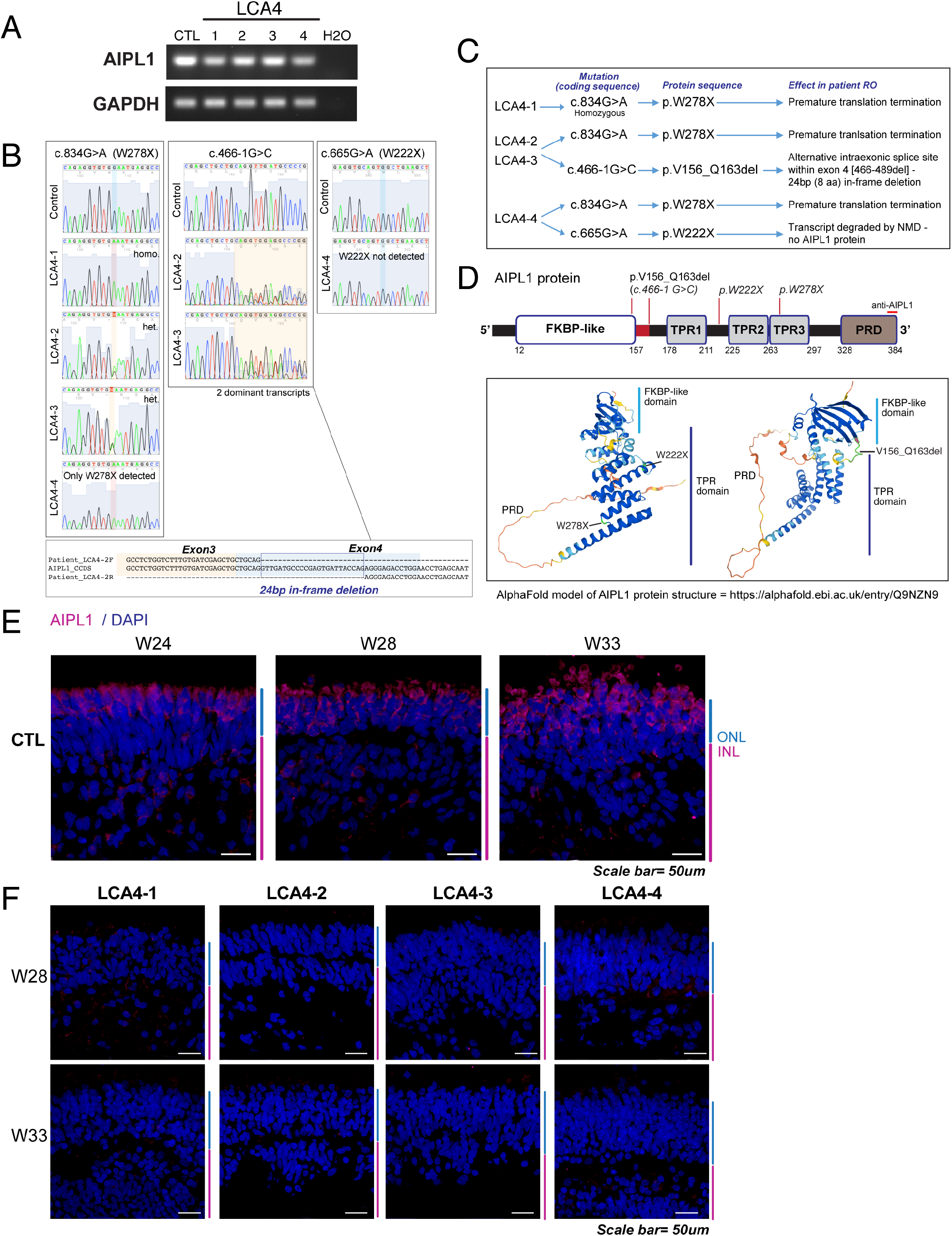
Analysis of AIPL1 transcript and protein in LCA4 ROs. A) AIPL1 transcript was present in all LCA4 ROs; semi-quantitative PCR reveals the presence of a PCR product for all samples. B) Amplification and Sanger sequencing of regions containing the c.834G>A (p.W278X), c.466-1G>C and c.665G>A (p.W222X) mutations from LCA4 cDNA revealed that p.W278X transcript was detectable in all LCA4 samples, while the p.W222X transcript was undetectable in LCA4-4. c466-1G>C gives rise to a differentially spliced mRNA transcript that excludes 24bp of exon 4. C) Summary of the impact of the LCA4-1, LCA4-2, LCA4-3 and LCA4-4 mutations on AIPL1 transcript and protein. D) Schematic diagrams of the AIPL1 2D and 3D protein structure. The location of the epitope targeted by the human-specific anti-AIPL1 C-terminal antibody is shown in the linear structure. The LCA4 patient mutation sites are highlighted. The 3D structure of AIPL1 is an AlphaFold model: https://alphafold.ebi.ac.uk/entry/Q9NZN9. E) IF analysis of AIPL1 in CTL ROs (W24-W33 of development) with the C-terminal human specific AIPL1 antibody. AIPL1 was present in the cell body and IS regions of the PRs located in the ONL. DAPI staining is in blue. ONL and INL regions are highlighted at the side of images in blue and magenta. Scale bars = 50μm. F) AIPL1 protein is not detectable in LCA4 ROs with the C-terminal human specific AIPL1 antibody. IF analysis of AIPL1 in W28 and W33 RO sections. DAPI staining is in blue. ONL and INL regions are highlighted at the side of images in blue and magenta. Scale bars = 50μm.

In order to ascertain if LCA4 ROs produce detectable AIPL1 protein, IF was carried out on W28 and W33 LCA4 RO sections with an AIPL1 polyclonal antibody targeted to the human-specific C-terminus of the protein.^6^ AIPL1 protein was first observed in CTL iPSC-RO at approximately W17 (data not shown) and gradually increased in intensity as the ROs matured (Figure 3E). AIPL1 protein in the LCA4 ROs was not detectable at any timepoint (Figure 3F). As the p.W278X mutation leads to premature translation termination and C-terminal truncation of AIPL1, the missing antibody epitope may account for the lack of detectable p.W278X expression in ROs from LCA4-1 (p.W278X homozygous) and LCA4-4 (compound heterozygous for p.W278X and p.W222X (transcript subject to NMD)). The lack of detectable AIPL1 protein in LCA4-2/-3 ROs (c.834G>A, p.W278X; c.466-1G>C) suggests that the c.466-1G>C transcript likely results in an AIPL1 protein species (p.V156_Q163del) that undergoes rapid degradation. The lack of detectable levels of AIPL1 in all LCA4 ROs was confirmed with a second well-characterised anti-AIPL1 antibody raised against recombinant purified full length human AIPL1 (Supplementary Figure 3C).^51^ Therefore, there was no detectable AIPL1 in any of the LCA4 ROs. Overall, this suggests that the p.W222X product is not detected as a result of NMD of the transcript, whereas p.W278X and p.V156_Q163del, the respective transcripts of which were expressed in the patient ROs, likely misfold and are rapidly degraded.

### LCA4 ROs recapitulate the key molecular features of the disease *in vitro*

Concomitant with the established role of AIPL1 in the post-translational stabilisation and correct assembly of the cGMP PDE6 subunits in rods and cones,^10^ cGMP PDE6 subunits have been shown to be greatly diminished in various AIPL1 loss of function animal models.^16–18,52,53^ To examine this in the iPSC-RO model, W28 LCA4 RO sections were immunostained for the rod cGMP PDE6 subunits, PDE6α and PDE6β. Both PDE6α and PDE6β were localised to the presumptive rod PR outer segments in the CTL RO, but were completely absent in the LCA4 RO PRs (Figure 4A). *PDE6A* and *PDE6B* transcripts were expressed in a similar temporal pattern in CTL and LCA4 RO (*see* Figure 2C) and were detectable in all W28 RO samples (Figure 4B), indicating that the loss of PDE6 protein occurs post-transcriptionally, as expected. In contrast to rod PDE6 proteins, examination of rhodopsin (RHO) and cone opsin (OPN1LW/MW, OPN1SW) localisation showed that the pattern of distribution and morphology of the PR cell populations do not appear to be different in LCA4 ROs compared to CTL ROs (Figure 4C).

**Figure 4:**
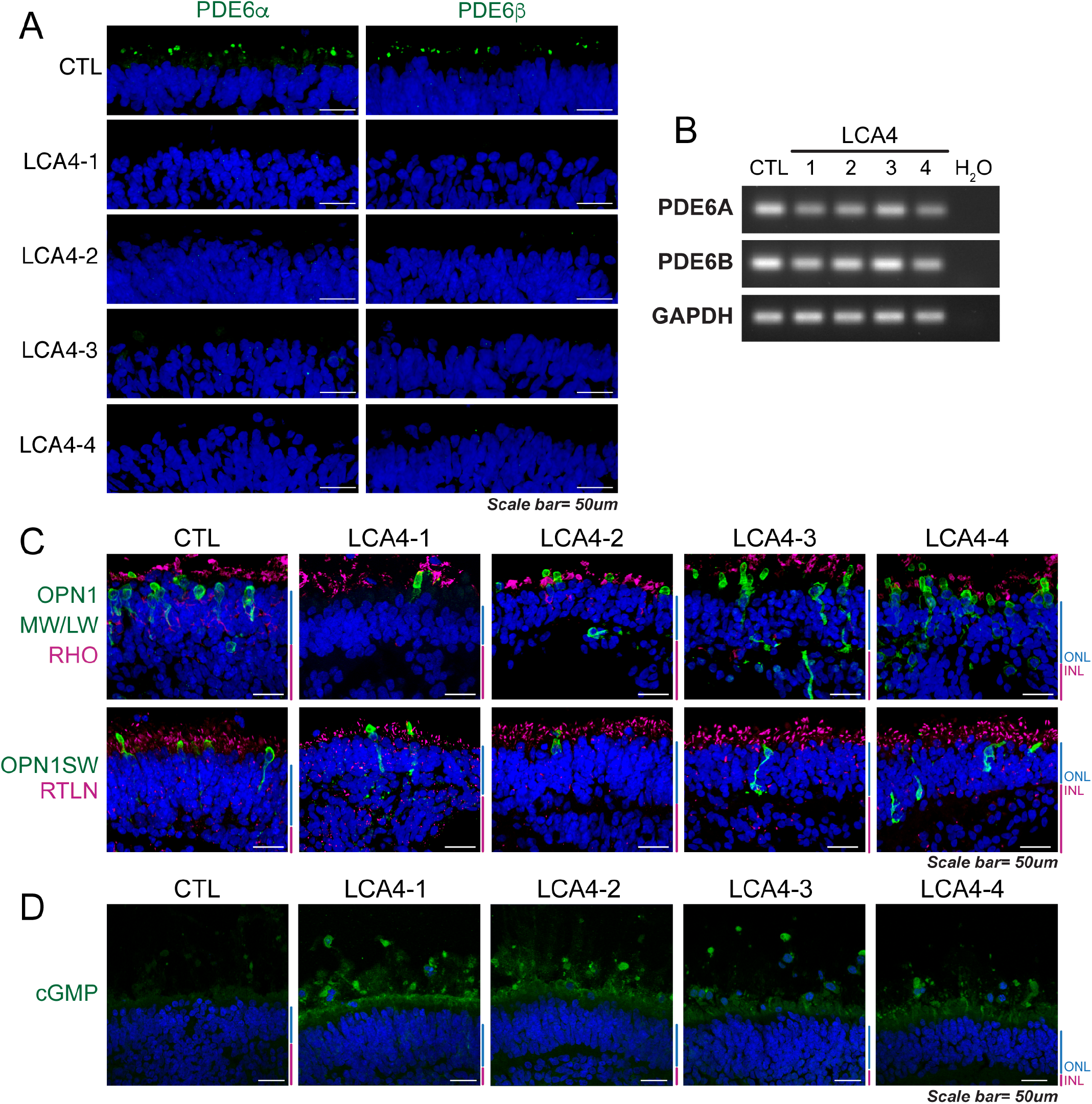
Photoreceptors in LCA4 ROs lacked detectable expression of rod cGMP PDE6α/PDE6β protein and displayed elevated levels of cGMP. A) IF analysis of rod cGMP PDE6αand PDE6βlevels. Images show the ONL region of W28 RO sections. PDE6α and PDE6β are normally observed in the OS regions of PRs (CTL panels), but the detection of PDE6α and PDE6β was absent in all LCA4 ROs. DAPI staining is in blue. Scale bars = 50μm. B) *PDE6A* and *PDE6B* transcripts were present in LCA4 ROs and were comparable to CTL levels, indicating post-translational mechanisms for loss of PDE6α and PDE6β protein. Semi-quantitative PCR, with housekeeping gene GAPDH for comparison. C) Other PR-lineage markers were unaffected in LCA4 ROs. IF analysis of W33 RO sections for Opsins (OPN1MW/LW, OPN1SW), Rhodopsin (RHO), and Rootletin (RTLN) revealed a similar pattern of expression in CTL and LCA4 ROs. DAPI staining is in blue. ONL and INL regions highlighted at the side of images in blue and magenta. Scale bars = 50μm. D) cGMP levels were elevated in LCA4 ROs; IF analysis of cGMP in W28 RO sections showed that CTL ROs have a low background level of cGMP positivity, in contrast to LCA4 ROs (cGMP positivity was noticeably bright in the IS/OS regions of PRs). DAPI staining is in blue. ONL and INL regions are highlighted at the side of images in blue and magenta. Scale bars = 50μm.

The PDE6 complexes play a crucial role in the hydrolysis of cGMP during the phototransduction cascade in PR cells. In order to ascertain if cGMP levels were raised in LCA4 ROs compared to CTL, W28 sections were immunostained for cGMP. CTL ROs did not exhibit a high background level of cGMP and no clear cGMP positive cells were observed. In contrast to this, the IS/OS regions of LCA4 RO PRs were clearly positive for cGMP (Figure 4D). The loss of PDE6 subunit proteins and the elevation of cGMP levels in LCA4 ROs phenocopies the molecular characteristics of LCA4, is in broad agreement with AIPL1 loss-of-function experimental models, and confirms the suitability of the model for the study of the disease *in vitro*.

### PTC124 treatment rescued full length AIPL1 in LCA4 ROs homozygous for the p.W278X stop mutation

Our data confirm that the position of the c.834G>A, p.W278X mutation enables the transcript to largely escape NMD and is therefore a potentially suitable target for TRID therapy. TRID molecules allow for the incorporation of cognate/near-cognate amino acids at PTC sites and the translation of full-length protein via a mechanism involving tight ribosomal binding.^54^ However there is evidence that PTC124 acts through a different mechanism whereupon it inhibits release factor (eRF1/eRF3) activity at the ribosomal complex to promote readthrough of PTCs (Figure 5A).^33^

**Figure 5:**
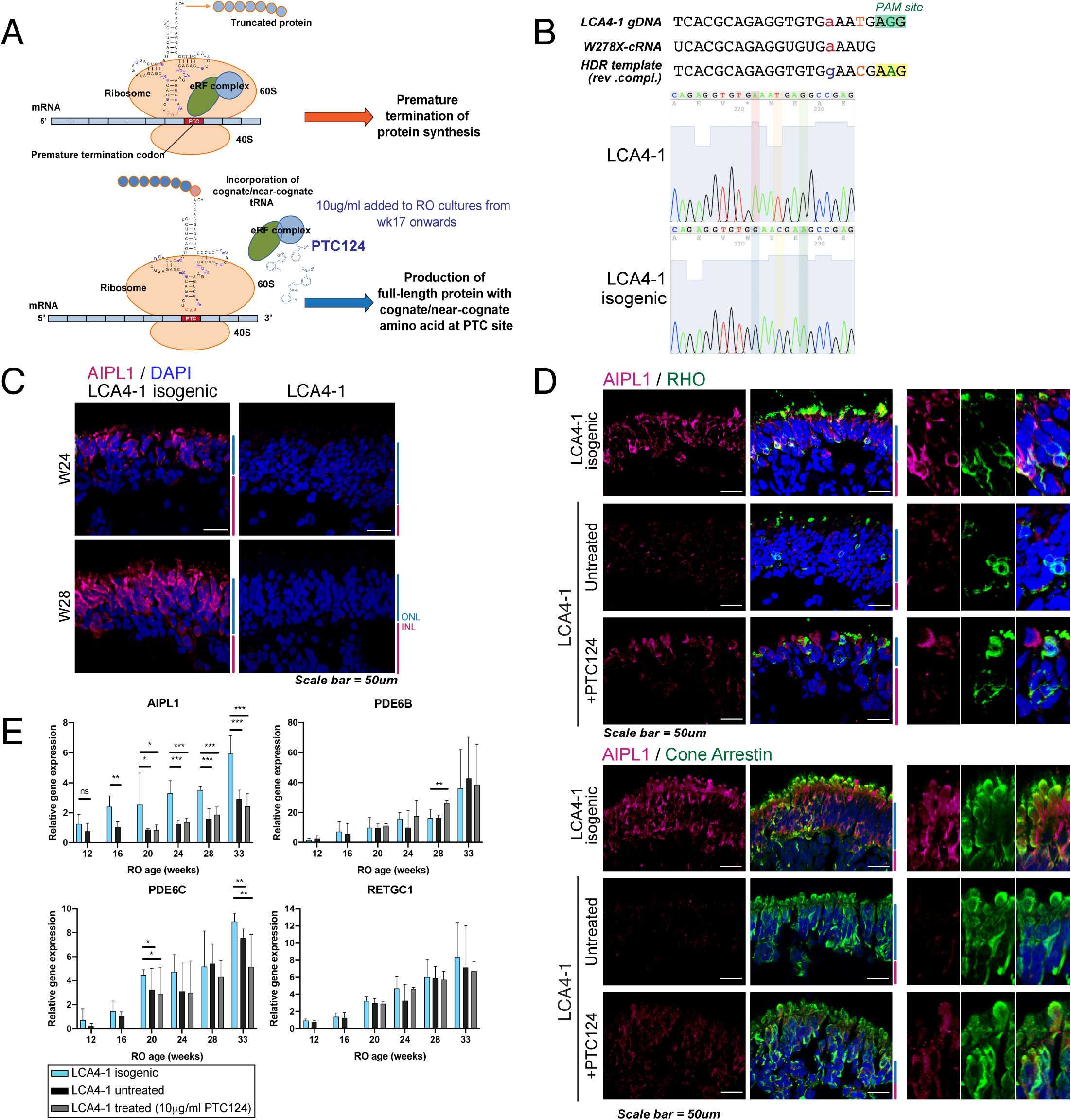
CRISPR-Cas9 repair of p.W278X in LCA4-1 iPSCs restored AIPL1 protein levels, and PTC124 treatment was able to partially rescue AIPL1 protein levels in LCA4-1 patient ROs. A) Schematic diagram detailing the proposed mechanism of action of PTC124 in driving translation readthrough of premature termination codons (PTCs). B) Schematic diagram detailing the CRISPR-Cas9 HDR correction strategy of the c.834G>A, p.W278X mutation. 2 additional changes (synonymous) were introduced in the HDR template to remove the PAM site and to prevent re-editing of the locus. Sequence chromatograms of unedited LCA4-1 compared to LCA4-1 isogenic repair cells. C) IF analysis of LCA4-1 isogenic ROs (W24, W28) demonstrated restoration of AIPL1 protein to the ONL PR cells, as observed in CTL ROs. Unedited LCA4-1 RO shown for comparison. DAPI staining is in blue. ONL and INL regions are highlighted at the side of images in blue and magenta. Scale bars = 50μm. D) AIPL1 levels were partially restored in both rod and cone lineage cells in PTC124-treated LCA4-1 ROs. IF analysis of ROs for AIPL1, RHO and Cone arrestin expression showed that AIPL1 was expressed in both RHO+ and Cone arrestin+ cells in PTC124-treated ROs. DAPI staining is in blue. ONL and INL regions highlighted at the side of images in blue and magenta. Scale bars = 50μm. E) Quantitative PCR analysis of *AIPL1*, *PDE6B* (rod PDE6), *PDE6C* (cone PDE6) and *RETGC1* levels in LCA4-1 isogenic, LCA4-1 untreated and treated (10ug/ml PTC124) ROs (W12-W33, 3 biological replicates per timepoint). While *PDE6B*, *PDE6C* and *RETGC1* levels were broadly similar for all RO types, *AIPL1* transcript levels were approximately halved in LCA4 (both untreated and PTC124-treated) ROs at all timepoints, compared to LCA4-1 isogenic ROs. Gene expression levels normalised to PR marker, *CRX*. Levels of *CRX* normalised to *ACTB* are shown in Supplementary Figure 6D. Values = mean +/− SD. (* ≤0.05 significance, ** ≤0.01 significance).

In order to generate optimal controls for the investigation of the effect of PTC124 on LCA4-1 ROs homozygous for the c.834G>A, p.W278X nonsense mutation, CRISPR-Cas9 homology directed repair (HDR) was carried out to generate isogenic repair lines (Figure 5B). Of the 2 sets of sgRNA and ssODN templates tested, one combination was efficient at triggering CRISPR-Cas9 HDR of the p.W278X locus (Supplementary Table 4) and was used to generate isogenic LCA4-1 iPSC lines, of which 2 were further characterised with regards to the expression of pluripotency markers and trilineage differentiation potential (Supplementary Figure 4). The isogenic lines were screened at 10 different genomic loci identified as potential sites for off-target editing by the sgRNA:SpCas9, and no off-target editing was observed (Supplementary Table 5). In ROs derived from the LCA4-1 isogenic iPSC, AIPL1 was detected specifically in the PR cells localised to the ONL (Figure 5C), similar to observations seen in CTL ROs. The CRISPR:Cas9 HDR repair therefore restored AIPL1 protein expression.

A pilot study short-term (2W) PTC124 dosage gradient (5-15ug/ml) of LCA4 ROs revealed that PTC124 could increase AIPL1 immunoreactivity, and the range of 10-12.5ug/ml was the most efficient at driving AIPL1 p.W278X readthrough in ROs (Supplementary Figure 5). PTC124 (10ug/ml) was added to LCA4 RO cultures from W17 onwards, as this is the timepoint at which AIPL1 protein expression was first observed in developing CTL ROs. Drug treatment was maintained throughout the remainder of the culture period. IF staining was carried out on untreated and PTC124 treated LCA4-1 ROs and isogenic CTLs with the AIPL1 C-terminal antibody to detect readthrough of full-length AIPL1 from the p.W278X transcript. IF staining of AIPL1 with RHO and cone arrestin showed that both markers colocalise with subsets of AIPL1 positive cells in the LCA4-1 isogenic ROs, demonstrating that AIPL1 is present in both rod and cone-lineage PRs (Figure 5D, W24 RO), as previously shown in developing human retina.^7^ In LCA4-1 ROs treated with 10ug/ml PTC124, low levels of restoration of AIPL1 immunoreactivity was observed compared to the isogenic CTL, and RHO+ AIPL1+ and CARR+ AIPL1+ cells were present in the treated ROs (Figure 5D, W24 RO). PTC124 is therefore able to promote readthrough of the AIPL1 PTC in both types of PR cells. PR markers (transducin, OPN1SW, RHO) were similar in LCA4-1 isogenic and LCA4-1 untreated and treated RO (Supplementary Figure 6A), and the level of cell death (TUNEL assay) was not increased in LCA4-1 RO or in the presence of 10ug/ml PTC124 (Supplementary Figure 6B). Therefore, longterm dosage with PTC124, whilst promoting the readthrough of full-length AIPL1 protein, did not appear to have deleterious, toxic effects on RO cell survival (W24-W33).

Gene expression analysis of W12-W33 CTL isogenic and LCA4-1 ROs (untreated/treated) for genes involved in early retinal development (*VSX2*), photoreceptor function (*RCVN*, *RHO*, *OPN1SW* and *OPN1MW/LW*) and non-PR retinal markers (*PKCA*, *PROX1*, *CRALBP*) indicated that these genes were regulated in a similar temporal pattern in the isogenic CTL and LCA4-1 ROs (untreated/treated) (Supplementary Figure 6C). Quantitative gene expression analysis of isogenic and LCA4-1 (untreated/treated) ROs at different timepoints of RO development surprisingly revealed that LCA4-1 ROs consistently express approximately half the amount of AIPL1 transcript, compared to the isogenic ROs (Figure 5E). This was an unexpected finding given that the AIPL1 p.W278X transcript is not expected to undergo NMD. Levels of the transcript were not changed in PTC124-treated ROs compared to untreated, and therefore PTC124 appeared to have no effect on AIPL1 p.W278X transcript stability in the ROs. In contrast to the changes in *AIPL1* levels, expression levels of rod *PDE6B*, cone *PDE6C* and *RETGC1* (the latter has been shown to be downregulated at the protein level in an *Nrl*−/− *Aipl1*−/− all-cone retinal murine model^52^) were comparatively similar in the isogenic and LCA4 ROs (untreated/treated) (Figure 5E). Transcript levels of the photoreceptor marker CRX were also comparable between all sample types (Supplementary Figure 6D). Overall, the findings indicate that PTC124 treatment, whilst driving readthrough of the AIPL1 PTC, does not appear to impact cell survival or gene expression in developing ROs.

### PTC124 treatment rescued rod PDE6 in p.W278X homozygous LCA4 ROs

As PTC124 was able to rescue low levels of full-length AIPL1 in LCA4-1 RO compared to the isogenic CTL, we investigated the downstream impact on the recovery of rod PDE6β and cGMP levels. The isogenic repair of c.834G>A, p.W278X in LCA4-1 RO not only restored the expression and correct localisation of AIPL1, but also that of PDE6β which localised to the presumptive rod PR outer segments (Figure 6A). LCA4-1 PTC124 treated ROs displayed rescue of PDE6β in a small subset of PRs compared to the isogenic CTL, with the protein correctly localised to the presumptive OS region of the cells (Figure 6A). Cells that displayed rescue of PDE6β were, however, relatively rare compared to the LCA4-1 isogenic CTL.

**Figure 6:**
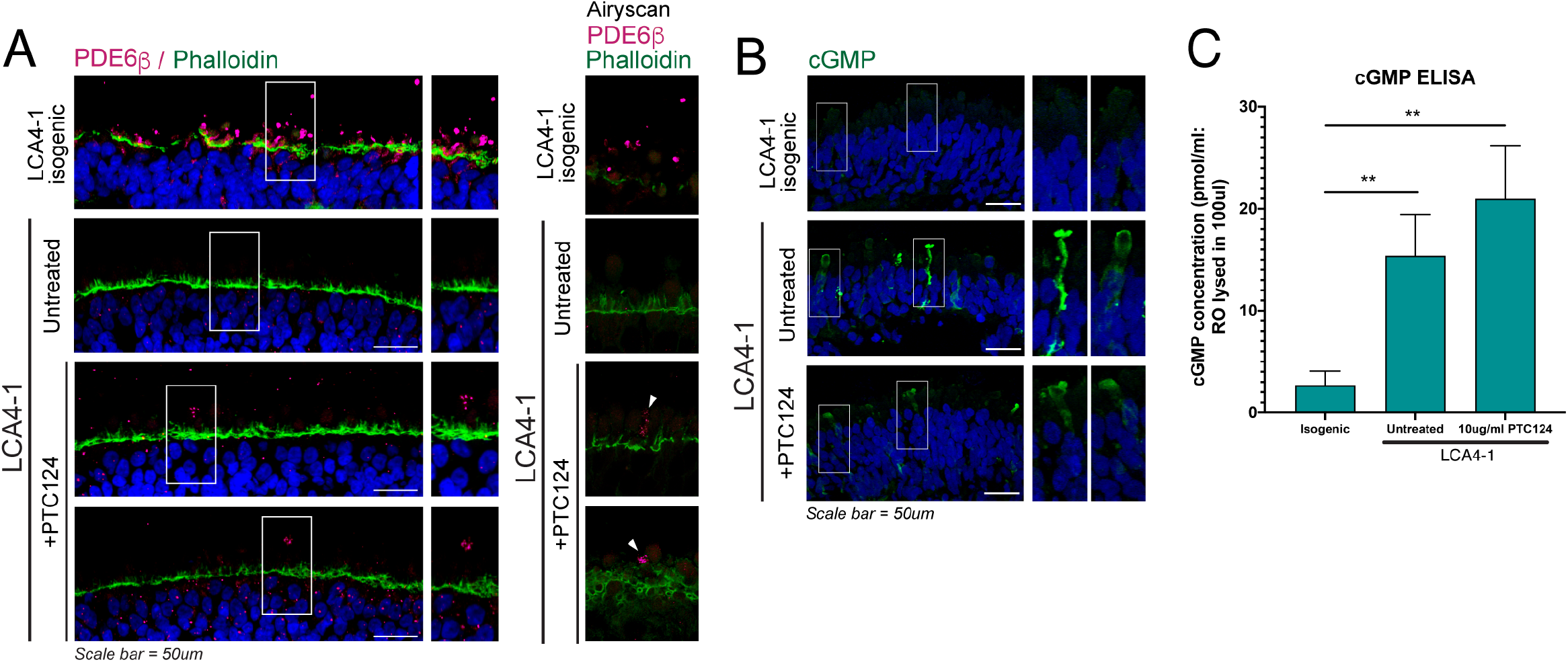
PTC124 treatment was able to restore low-level PDE6βto LCA4-1 ROs, but had no impact on cGMP levels. A) IF analysis of ROs for PDE6β. Images show the ONL region of W28 ROs. PDE6β was abundantly expressed in the OS regions of PRs in LCA4-1 isogenic ROs. Rescue of PDE6βwas observed in PTC124-treated LCA4-1 ROs; a small number of PR cells displayed PDE6β staining localised to the IS/OS region (white arrowheads, Airyscan images). DAPI staining is in blue. Scale bars = 50μm. B) cGMP IF analysis showed that LCA4-1 isogenically repaired ROs displayed a low background level of cGMP. Images show the ONL region of W28 ROs. Bright cGMP+ PR cells were present in both LCA4-1 untreated and PTC124-treated ROs. DAPI staining is in blue. Scale bars = 50μm. C) cGMP ELISA analysis of whole W33 ROs confirmed the elevation of cGMP levels in LCA4-1 ROs; treatment with PTC124 had no effect on cGMP levels. 3 biological replicates; mean +/− SD. **: ≤0.01 significance level).

As expected, similar to CTL ROs, cGMP levels were undetectable in PR cells in LCA4-1 isogenic ROs (Figure 6B). As previously observed, the level of cGMP was elevated in LCA4-1 RO cells and a number of high cGMP+ cells were visible, with the cGMP distributed throughout the length of the PR cell structure. PTC124 treatment, however, had no discernible effect on the pattern or intensity of cGMP distribution (Figure 6B). This finding was confirmed by cGMP ELISA results with whole organoid material, which revealed low levels of detectable cGMP in isogenic CTL organoids and the significant elevation of cGMP in the LCA4-1 patient RO. There was no significant change in cGMP levels in PTC124-treated compared to untreated ROs (Figure 6C). Therefore the readthrough of low levels of full-length AIPL1 and the restoration of PDE6 in a limited number of PRs was not sufficient to reduce cGMP levels in whole organoids.

### PTC124 induced readthrough of AIPL1 p.W278X and rescued full length AIPL1 in LCA4 ROs compound heterozygous for the stop mutation

As the c.834G>A, p.W278X mutation is prevalent in LCA4 patients compound heterozygous for this allele, we also tested PTC124-mediated readthrough in LCA4-2 ROs that are compound heterozygous for c.834G>A, p.W278X and the c.466-1G>C splice mutation. ROs between W20-W33 were collected to study the effect of PTC124 treatment on AIPL1 protein levels. The IF results with the AIP1L C-terminal antibody indicated that there was partial rescue of AIPL1 levels at all the timepoints studied, and thus PTC124 is indeed also able to drive readthrough of the AIPL1 p.W278X transcript in LCA4 p.W278X compound heterozygous ROs (Figure 7A: 10x magnification images (W28 RO), Figure 7B: 40x of ONL region (W20-33 RO)). The rescued AIPL1 protein localised to the PR cells in the ONL of organoids, as expected, and had a similar intracellular cytoplasmic pattern of distribution as seen in CTL RO cells. As seen in PTC124-treated LCA4-1 ROs, PTC124 treatment had no obvious effect on the morphology of retinal cell types, including rods and cones, or retinal tissue layers (Figure 7C).

**Figure 7:**
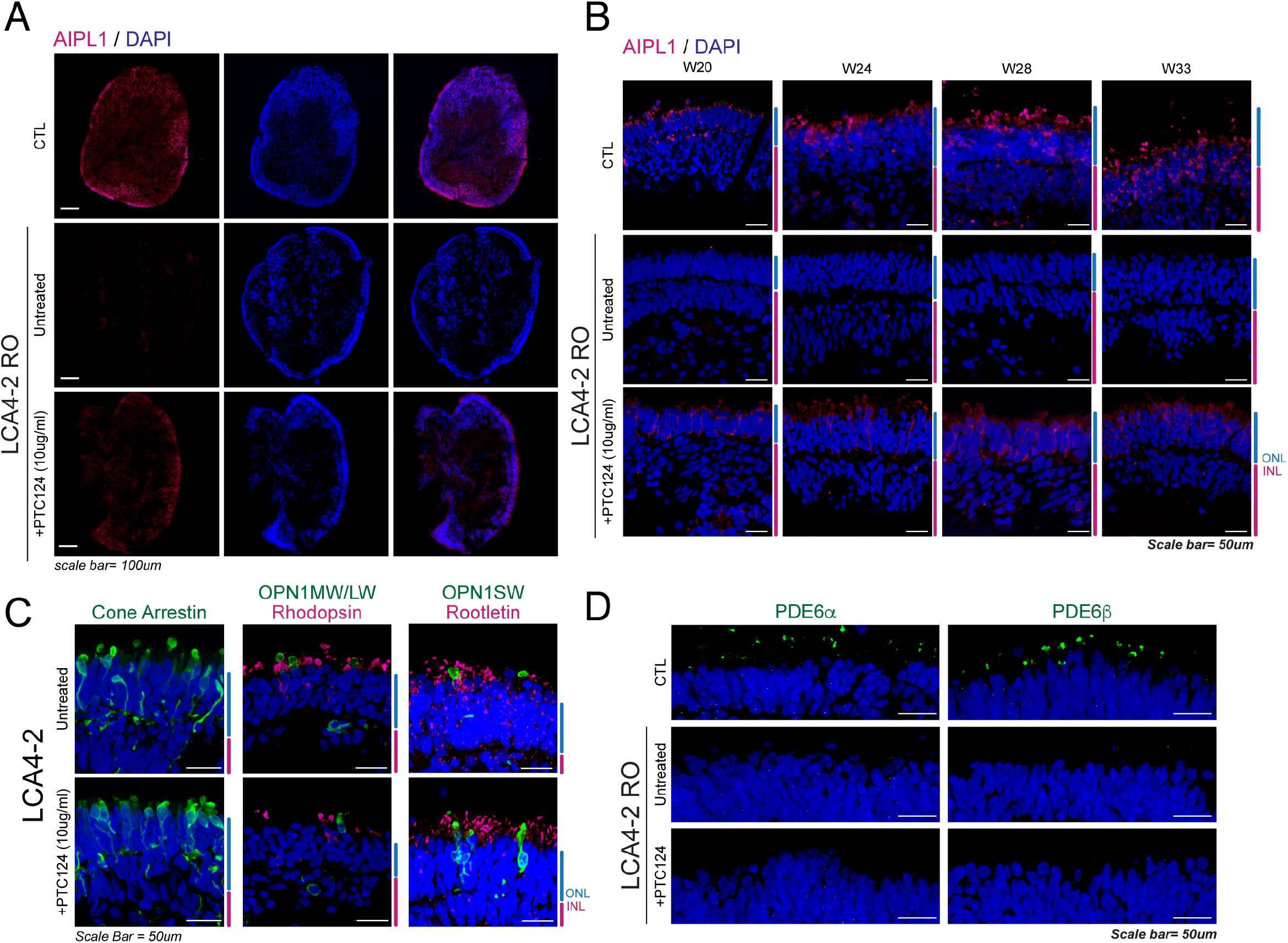
PTC124 translation readthrough treatment was able to partially rescue AIPL1 levels in LCA4 compound heterozygous ROs. A) AIPL1 IF of W28 CTL, LCA4-2 untreated and treated (10ug/ml PTC124) RO sections shown in whole organoids. AIPL1 levels were partially restored in LCA4-2 treated ROs. DAPI staining is in blue. Scale bars = 100μm. B) IF analysis of AIPL1 levels in W20-W33 CTL and LCA4-2 ROs (untreated / treated with 10ug/ml PTC124). Partial rescue of AIPL1 protein levels was observed in PTC124-treated ROs at all timepoints. DAPI staining is in blue. ONL and INL regions highlighted at the side of images in blue and magenta. Scale bars = 50μm. C) Treatment of LCA4 ROs with PTC124 (10ug/ml) had no effect on photoreceptor populations. IF for Cone Arrestin, OPN1MW/LW, Rhodopsin, OPN1SW and ciliary rootlet protein Rootletin in LCA4-2 untreated/treated ROs yielded similar results. DAPI staining is in blue. ONL and INL regions are highlighted at the side of images in blue and magenta. Scale bars = 50μm. D) IF analysis of PDE6α and PDE6β levels revealed a lack of rod cGMP PDE6 rescue in PTC124-treated LCA4-2 ROs. The images show the ONL region of W28 RO sections. DAPI staining is in blue. Scale bars = 50μm.

As PTC124 was able to rescue low levels of AIPL1 protein in treated LCA4-2 ROs, IF was conducted to ascertain if this was sufficient to partially rescue the rod PDE6 phenotype of LCA4-2 ROs, similar to observations in LCA4-1 ROs. Analysis of W28 sections for PDE6α and PDE6β protein demonstrated that in contrast to LCA4-1 ROs homozygous for the stop mutation, there was no observable elevation of PDE6 protein levels in the PTC124-treated ROs (Figure 7D).

## Discussion

In this study we have developed and characterized the first human AIPL1-LCA4 iPSC-retinal organoid (RO) system derived from renal epithelial cells. Moreover, we have developed an isogenic control through the homozygous repair of the c.834G>A, p.W278X mutation in the LCA4-1 patient line. This *in vitro* model constitutes a powerful tool to understand different mechanisms of pathogenesis of *AIPL1* variations. We have identified that the transcripts from the AIPL1 c.466-1G>C splice variation and c.834G>A, p.W278X nonsense variation result in AIPL1 protein species that undergo rapid degradation. In contrast the AIPL1 c.665G>A, W222X nonsense variation involves the production of a transcript that undergoes NMD.

We show that ROs from four LCA4 patients, sharing 3 different *AIPL1* genotypes, generated layers of cells that reflect those found in endogenous developing retinal tissue; photoreceptors in the ONL, bipolar, amacrine and horizontal cells in the INL and retinal ganglion cells underlying the INL. The LCA4 ROs were indistinguishable from control organoids in terms of size and ONL-INL lamination, and the photoreceptor cell population demonstrated no noticeable structural differences. There were also no overt signs of retinal cell death or degeneration in the LCA4 retinal organoids. The absence of functional AIPL1 protein in the LCA4 ROs therefore seems to have no bearing on the specification or the maturation of the photoreceptor cells (both rods and cones) or the other retinal cell lineages that are seen in the RO differentiation model. These findings are in agreement with a recently published model of LCA4 harbouring the homozygous AIPL1 mutation p.C89R, which showed normal retinal development and cell type specification and maintenance of the retinal cytoarchitecture.^55^ This is also consistent with loss of function animal models of LCA4 which similarly showed no gross abnormalities or cell death during retinal development^17,18,56^.

We moreover report the loss of PDE6α and PDE6β in the patient-derived ROs confirming that the patient ROs recapitulate the molecular features of LCA4. In addition to the expected loss of AIPL1 and rod cGMP PDE6 subunits, we show that cGMP levels are elevated in LCA4 photoreceptors. Similarly, cGMP levels have been reported as raised in early post-natal stage *Aipl1* knockout whole mouse retinas.^18^ This is in contrast, however, to the results from *Aipl1* hypomorphic whole mouse retinas, which reported decreased levels of cGMP.^17^ Moreover, cGMP levels were reported as decreased in an all-cone mouse model of *Aipl1* loss-of-function,^52^ and significantly elevated levels of cGMP were not detected in a zebrafish cone-specific *aipl1b* mutant.^53^ Interestingly, in both the all-cone mouse *Aipl1* knockout model and the zebrafish cone-specific *aipl1b* mutant, reduced levels of cone PDE6 were reported in addition to reduced mouse retinal membrane guanylate cyclase (RetGC1) and zebrafish cone-specific guanylate cyclase (zGc3), respectively, in cone photoreceptors. This has been suggested to point to mechanistic differences in rod and cone cell death caused by deficiencies in AIPL1 differentially affecting cGMP homeostasis. Indeed, higher cGMP levels are observed in rod PDE6 loss-of-function animal models^57–60^ and human retinal organoids,^61^ while the RetGC1/RetGC2 double knockout mouse exhibits a lack of cGMP.^62^

We did not observe increased apoptosis or changes to the photoreceptor cell population in LCA4 retinal organoids despite the increased cGMP levels. In many IRDs, increases to cGMP levels precede the degeneration of photoreceptor cells, and there is evidence that cell death is primarily driven by non-apoptotic pathways, with increases in calpain, histone deacetylase (HDAC) and poly-ADP-ribose-polymerase (PARP) activity detected and comparatively low-level activation of BAX, caspases, and other hallmarks of the apoptotic pathway.^63^ Whilst disturbances in calcium flux in cGMP-elevated conditions have a deleterious effect on photoreceptor survival,^64,65^ cGMP may also act via other pathways to negatively impact cell survival. The activation of the cGMP-dependent Protein Kinase G (PKG) pathway has been implicated in photoreceptor cell death^66^ and increased PR survival reported with PKG ablation in a mouse model that lacks cGMP gated channel *CNGb1* function.^65^ However, evidence of overt PR degeneration is lacking in the LCA4 RO, and PR degeneration proceeds after birth in both LCA4 patients and animal models, suggesting that the RO model may suitably recapitulate retinal development events but not the molecular events immediately preceding PR degeneration.

The use of patient-specific ROs derived from iPSC constitute a powerful tool to test new therapeutics for IRDs. Our data shows that the treatment of ROs homozygous or compound heterozygous for the c.834G>A, p.W278X mutation with the translation readthrough inducing drug PTC124 rescued full-length AIPL1 protein. This was sufficient to rescue low levels of PDE6 in p.W278X homozygous ROs, but not the increased cGMP levels, suggesting that whilst increased levels of full length AIPL1 protein were gained by the treatment, the restoration of wild type functional AIPL1 was not sufficient to restore PDE6 to the levels required to reduce cGMP levels. The efficiency of readthrough is dependent on a number of factors, which in addition to the TRID itself, include the amount of target transcript, the nucleotide context of the premature translation codon (PTC) and the features of the full-length protein arising from the readthrough event.

As expected, the efficiency of phenotypic rescue was greater in retinal organoids homozygous for c.834G>A, p.W278X due to the increased availability of the PTC-bearing transcript. The c.834G>A, p.W278X mutation is located 50 nucleotides downstream of the final exon 5-exon 6 junction, and the transcript is thus expected to be resistant to NMD. Indeed, the AIPL1 c.834G>A, p.W278X transcript was detected in all patient-derived retinal organoids, however we also observed that levels of *AIPL1* transcript were elevated in the LCA4-1 isogenic control ROs compared to uncorrected ROs, but the mechanism for why this might be the case has yet to be determined. A combined approach that both promotes readthrough and inhibits NMD could therefore potentially enhance the nonsense suppression efficiency of this PTC. Increasing the concentration of PTC124 in our RO model did not further enhance the readthrough level (data not shown) indicating a threshold for readthrough efficiency mediated by this drug in our system.

With respect to the nucleotide context, PTC124 selectively induces readthrough of PTC over natural stop codons and promotes translational readthrough of all three stop codons, with the highest efficiency for UGA followed by UAG and UAA.^67,68^ The resultant c.834G>A, p.W278X termination codon is UGA, the most amenable to PTC124 induced readthrough and therefore expected to yield the highest levels of readthrough. However, the efficiency of PTC124 is also influenced by the nucleotide context surrounding the PTC with the nucleotide immediately 3’ to the termination codon strongly implicated in readthrough efficiency and termination fidelity, wherein increased efficiency is favoured by a pyrimidine base, especially cytosine.^67,68^ This position is occupied by adenine at the c.834G>A, p.W278X locus, though readthrough efficiency cannot be predicted by the nucleotide context of the PTC alone. In terms of the protein context, the insertion of a near-cognate tRNA coding for tryptophan at the UGA PTC, which would reinstate the AIPL1 wild type sequence, is only one of several possibilities. The characterization of translational readthrough products from mammalian 293H cells treated with PTC124 revealed the predominant insertion of arginine (~69%), followed by tryptophan (~28%) and cysteine (~0.7%) at the UGA PTC.^31^ Therefore, less than a third of the rescued AIPL1 recovered in PTC124-treated retinal organoids may be wild type full length functional AIPL1 harbouring a reinstated tryptophan residue. *In silico* predictions of p.W278R and p.W278C predict that insertion of arginine or cysteine respectively are neutral or probably deleterious (Rhapsody),^69^ or that both are deleterious (PolyPhen2). Moreover, the W278 residue occupies an important structural position in the AIPL1 TPR domain, and the substitution of tryptophan with arginine or cysteine is likely to be poorly tolerated at this position. In summary, we showed that readthrough restores the production of full-length AIPL1, albeit at levels considerably reduced from wild type. Heterozygous carriers of disease-associated *AIPL1* variations are unaffected, suggesting that sub-threshold levels of readthrough combined with heterogeneity in the incorporation of the near-cognate amino acid during AIPL1 translation could explain the sub-therapeutic rescue and amelioration of the disease phenotype. Ng *et al*.^33^ showed that PTC124, which generally results in lower yields of readthrough product, potentiated G418-stimulated readthrough suggesting additivity of the combined action of these TRIDs. Therefore, the recovery of PDE6 in a small number of cells suggests that combinatorial therapies based on the orthogonal mechanisms of action of PTC124 and less toxic aminoglycoside derivatives aimed at further upregulating the readthrough of full length wild type AIPL1 could be beneficial.^33^

Overall, the consistent pathogenic features and disease phenotype of the AIPL1 associated-LCA4 RO model developed in this study render it ideal for the dissection of the molecular mechanisms that precede cell death in LCA4 retinas, and for evaluating the efficacy of new drugs or gene therapy products.

## Experimental Procedures

### Ethical Approval

To collect urine samples, parents or legal guardians of patients (who were all children under the age of 3 years) signed an informed consent in adherence with the Declaration of Helsinki and with approval from the North East - Newcastle & North Tyneside 2 Research Ethics Committee.

### Clinical Data and Patient Imaging

Medical notes and clinical images were reviewed. This included results of comprehensive ophthalmic clinical assessment, including dilated fundoscopy and age-appropriate visual acuity assessments, and electrophysiological testing. Optical coherence tomography (OCT) imaging was reviewed. OCT was acquired with handheld Bioptigen spectral domain OCT (Leica Microsystems, Research Triangle Park, NC, USA). Trans-foveal horizontal scan of a healthy adult was acquired using Heidelberg Spectralis OCT (Heidelberg Engineering, Heidelberg, Germany).

### Urine Collection, Cell Isolation and Expansion of Renal Epithelial Cells

Urine was collected from LCA patients with mutations in the *AIPL1* gene. Collection, isolation, and expansion of renal epithelial cells was performed similarly to described previously.^48,70^ Briefly, urine samples (volumes varying from 15mls to 80mls) were collected in sterile tubes and centrifuged at 400g for 10min at room temperature. After discarding the supernatant, pellets were washed with 10mls of PBS containing 500ng/ml ampothericin B and 100 U/ml penicillin/streptomycin and centrifuged again at 400g for 10min. Pellets were resuspended in 2mls of Primary Medium consisting of DMEM/Ham’s F-12 nutrient mix (1:1) (ThermoFisher Scientific), with 10% of fetal bovine serum (FBS), Renal Cell Growth Medium (REGM) SingleQuot kit supplements (Lonza), 2.5μg/ml amphotericin B, and 100 U/ml of penicillin/streptomycin. The cells were seeded into one well of a 12-well plate coated with 0.1% gelatin. One ml of Primary Medium was added to the well at 24h, 48 and 72h without removing any media. Renal epithelial cells routinely appear within the first 3-5 days of culture. 96h post seeding, most of the medium was removed and replaced with Proliferation Medium, consisting of (1:1 mixture of Renal Cell Growth Medium (REBM) medium supplemented with REGM SingleQuots (Lonza) and DMEM high glucose (ThermoFisher Scientific) supplemented with 10% FBS, 1% GlutaMAX, 1% non-essential amino acids (NEAA), 100 U/ml penicillin/streptomycin. Subsequently, half of the culture medium was changed every day. Once cell density reached 90% (between 14–28 days after urine collection), cells were split 1:4 using 0.25% Trypsin-EDTA, (ThermoFisher Scientific) and expanded for a maximum of four passages.

### Reprogramming, Culture, and Characterization of iPSCs

Renal epithelial cells passaged fewer than 3 times were used for iPSC generation as described previously.^71^ Briefly, 5 × 10^5^ cells at early passages were trypsinized (0.25% Trypsin-EDTA, ThermoFisher Scientific) and electroporated with integration-free episomal plasmids pCXLE-hOCT3/4-shp53-F, pCXLE-hUL, and pCXLE-hSK (Addgene) and miRNA 302/367 plasmid (Gift from Dr J. A.Thomson, Regenerative Biology, Morgridge Institute for Research, Madison, Wisconsin, USA) using the Amaxa™ Basic Nucleofector™ Kit for primary mammalian epithelial cells, program T-020 (Lonza). Electroporated cells were plated onto geltrex-coated 12 well plates and cultured in E8 media (Gibco), which was changed every day. The iPSC colonies were picked around day 14, expanded in E8 on geltrex-coated 6 well plates and routinely passaged using Versene (Gibco). Pluripotency of the isolated iPSC lines was confirmed by immunofluorescence (IF), using iPSC-specific antibodies (Supplementary Table 1). iPSC cultures were grown in 8-well permanox chamber slides (ThermoFisher Scientific) and fixed in 4% PFA:PBS for 15 minutes at RT prior to IF. Genomic DNA was extracted for amplification of the *AIPL1* gene and PCR products were Sanger sequenced to confirm the presence of *AIPL1* mutations in the different patient lines (see Supplementary Table 2 for primer sequences). To confirm trilineage differentiation potential, iPSCs were differentiated into the three germ layers using the StemDiff™ Trilineage Differentiation Kit (STEMCELL Technologies) according to manufacturer instructions, and the resultant tissues analysed for the expression of key germ layer markers by real time PCR (Supplementary Table 3).

### CRISPR-Cas9 Homology-Directed Repair (HDR) of AIPL1 c.834G>A, p.W278X allele

To correct the c.834G>A, p.W278X mutation, 20bp guide RNAs (gRNAs) were designed around the locus (NGG PAM) using Benchling software. 127bp single stranded oligo deoxynucleotide (ssODN) repair templates (antisense to the target strand) were designed for homology directed repair (HDR) of the mutation site, with additional synonymous changes introduced to remove the PAM site/prevent the reannealing and re-editing of the locus. See Supplementary Table 4 for sequences. CRISPR RNA (crRNA) and ssODN templates (high quality ultramer oligos with phosphorothioate (PS) modification of the two 5’ and 3’ nucleotides) were obtained from IDT. iPSC cultures were grown in Stemflex (Gibco) supplemented with 10uM ROCK inhibitor Y-27632 (StemCell Technologies) for 2 hours prior to single cell dissociation with TrypLE (Gibco). 2 × 10^5^ cells/sample were nucleofected with 130pmol of crRNA:tracrRNA duplex (Cas9 nuclease V3 tracrRNA) (IDT) complexed with 125pmol of Alt-R Cas9 V3 enzyme (61 uM) (IDT), 200pmol ssODN template and 120pmol Alt-R Electroporation enhancer (IDT), using a P3 Primary Cell 4D-Nucleofector X Kit S (Lonza). 2 nucleofections were tested for each gRNA:ssODN combination, either with or without HDR enhancer (IDT). After 10min at RT post-nucleofection, cells were plated onto rhLaminin-521 (ThermoFisher Scientific) coated wells (24-well plates) in Stemflex + ROCKinhibitor, and cultured thereafter in Stemflex until characterisation and single cell cloning to isolate correctly edited iPSC clones. 3 correctly edited clones (homozygous correction of c.834G>A, p.W278X) were established from LCA4-1 iPSC using sgRNA 1 and ssODN repair template 1 (**Supplementary Table 4**), of which 2 (LCA4-1 isogenic 1 and 2) were further characterised with respect to pluripotency markers and trilineage potential (**Supplementary Figure 4**). Potential genomic off-target sites were identified using Cas-OFFinder (http://www.rgenome.net/cas-offinder/)^72^ and Offspotter (https://cm.jefferson.edu/Off-Spotter/). Primers were designed to amplify the top 9 regions and products from LCA4-1 and the 2 isogenic lines were subject to Sanger sequencing – no changes were detected in the isogenic lines (see **Supplementary Table 5** for further information and primer sequences). LCA4-1 isogenic line 2 was used for all experiments.

### Differentiation of Retinal Organoids (ROs)

Directed differentiation of iPSCs into 3D ROs was based on the protocol described previously in Gonzalez-Cordero *et al*. 2017.^28^ iPSCs were dissociated with Versene (Gibco), clumps were collected, washed twice with PBS, resuspended in Essential 8™ (Gibco) medium, and seeded onto 6 well plates coated with Geltrex (Gibco). iPSC colonies were grown until 90-95% confluent, then Essential 6™ media (Gibco) was added for 2 days (Day 1 and Day 2 of differentiation) followed by a neural induction period in Neural Induction Media (Advanced DMEM/F12 (1:1, Gibco), 1% non-essential amino acids (Gibco, NEAA), 1% N2 Supplement (Gibco), 1% GlutaMAX (Gibco) and 100 U/ml penicillin/streptomycin (Gibco)). Some cultures were supplemented with human recombinant bone morphogenetic protein 4 (rhBMP4) to improve neural induction efficiency; on Day 6, media was supplemented with 1.5nM BMP4 (R&D Systems), and half media changes were carried out every other day until Day 16 to dilute the rhBMP4. Around week (W) 6, neuro-retinal vesicles (NRVs) were manually excised using 21G needles/scalpel blades and grown in low-binding 96 well plates (96 well Nunc Sphera Round Bottom Plates, ThermoFisher Scientific) in Retinal Differentiation Media (DMEM/F12 nutrient mix (3:1 ratio, Gibco), 10% fetal bovine serum (FBS, Gibco), 2% B27 supplement (without vitamin A), 100uM taurine, 2mM GlutaMAX and 100U/ml penicillin/streptomycin), with media changes every 2 days. At W10, cultures were supplemented with 1uM retinoic acid (RA), and ROs were transferred into low binding 24-well plates. At W12, the cultures were supplemented with 1% N2 and the RA concentration was reduced to 0.5uM. At W14 (D100), B27 and RA were removed from the medium.

### Translational Readthrough Treatment

The translational readthrough inducing drug Ataluren (3-[5-(2-fluorophenyl)-1,2,4-oxadiazol-3-yl]-benzoic acid), also called Translarna™ or PTC124, was purchased from Selleckchem. Aliquots in DMSO (10mg/mL) were stored at −20°C. Translational readthrough treatment of retinal organoids with PTC124 started at W17. Fresh PTC124 was diluted in Retinal Maturation Media at a final concentration of 5-25ug/ml and added to the retinal organoids every other day when changing media.

### Immunofluorescence and Imaging

Retinal organoids were fixed in 4% PFA, 5% sucrose in PBS for 30min at 4°C and dehydrated in 6.25%, 12.5% and 25% sucrose:PBS (1hr incubations at 4°C, ROs left in 25% sucrose overnight). Retinal organoids were embedded in OCT compound (Tissue-Tek) and stored at - 80°C. 7um cryosections were mounted on SuperFrost Plus™ slides (Thermo Scientific). For ICC, slides were incubated in 10% donkey serum (Sigma Aldrich) or FBS (Gibco), 0.01% Triton-X (Sigma Aldrich) in PBS for 1hr at room temperature (RT), before a 1hr primary antibody incubation (Supplementary Table 1: List of primary antibodies). Slides were washed 3 times with PBS, incubated with 1:1000 species-specific secondary antibody (Supplementary Table 1: List of secondary antibodies) for 1hr, washed 3 times and incubated with 4’,6-Diamidino-2-phenylindole (DAPI; 2mg/ml) (Invitrogen) in PBS for 5 minutes. Slides were dried at RT and mounted in Fluorescence Mounting Media (Dako). TUNEL staining was carried out on PFA-fixed sections using the *In Situ* Cell Death Detection Kit, Fluorescein (Roche) according to manufacturer’s instructions. All images were acquired using a Zeiss LSM700 and LSM710 laser-scanning confocal microscope. Images were exported from Zen 2009 (Zeiss) software and prepared using Adobe Photoshop, Image J and Adobe Illustrator CS6.

### RNA Extraction and Quantitative PCR (qPCR)

RNA from iPSC and ROs was extracted using RNeasy Micro Kit (Qiagen). cDNA synthesis was performed using the Tetro cDNA synthesis kit (Bioline). 2x GoTaq Green master mix (Promega) was used for DNA amplification by PCR with standard cycling conditions for semi-quantitative PCRs. Real time PCR reactions were set up with 2x LabTaq Green Hi Rox Master Mix (Labtech) and validated primers (at a concentration of 0.25pmol/ul), and run on an Applied Biosystems QuantStudio ™ 6 Flex real-time PCR system. Primer sequences used for semi-quantitative/qPCR are detailed in Supplementary Table 3. Gene expression levels were calculated using the ΔΔCt method; RO markers were normalised against CRX and also betaactin. CRX was chosen as a consistently expressed PR-specific reference gene to negate differences in RO size/cellular make-up between samples.

### Transmission Electron Microscopy

ROs were processed for TEM analysis and imaged as previously described in Lane *et al*, 2020.^73^

### cGMP ELISA

96-well Cyclic GMP ELISA kits (Cayman Chemicals) were used according to manufacturer’s instructions. To prepare ROs for ELISA, ROs were washed with PBS and then incubated in 100ul of 0.1M HCl for 20 minutes at RT, before mechanical homogenisation with a pipette. The samples were centrifuged at 1000g for 10 minutes, and supernatants decanted into clean Eppendorf tubes. 200ul of ELISA buffer was added to each sample. 50ul was used per ELISA well (3 technical replicates per sample); samples were not acetylated for the analyses. Absorbance was measured at a wavelength of 420nm.

### Statistical Analysis

For real time PCR analysis of gene expression levels and the cGMP ELISA analyses, the values from 3 biological replicates per sample/RO type were taken into account for the calculations of group averages and standard deviations (SD). Pairwise comparisons were carried out using 2-tailed Student’s t-tests (* ≤0.05 significance, ** ≤0.01 significance, annotated on the relevant graphs).

## Acknowledgements

This work was funded by the Medical Research Council (to J.v.d.S (PI) and Co-I M.E.C, P.C., J.B., M.M). We would like to thank Dr Anai Gonzalez-Cordero (Faculty of Medicine and Health, University of Sydney, Australia) and Arifa Naeem (MeiraGTx, UK) for advice on the retinal organoid differentiation protocol. We would also like to thank Professor Viswanathan Ramamurthy (Departments of Ophthalmology, Biochemistry and Pharmaceutical and Pharmacological Sciences, West Virginia University, USA) for the AIPL1 antibody.^51^ We are extremely grateful to the patients and their families for supporting this research.

## Author Contributions

A.L., A.S.-R., P.R.L.P and H.S. performed the experiments and/or analysed the data. M.G., and A.K. performed clinical studies and data analysis. A.F.C., P.J.C., M.M., J.B., and M.E.C. provided materials, laboratory samples or patients for the research. A.L., A.S.-R. and J.v.d.S. conceived the hypothesis and designed the experiments, and drafted the manuscript. All authors edited the draft manuscript.

## Figure Legends

**Supplementary Figure 1:** Optical coherence tomography (OCT) images from LCA4 patients.

A) Patient LCA4-1 (p.W278X homozygote) - OCT image at 3.2 years of age.

B/C) Patients LCA4-2 and LCA4-3 respectively (monozygotic twins; c.834GA, p.W278X; c.466-1G>C compound heterozygotes) - OCT images at 2 years old.

D) OCT of a healthy control patient (adult, male, 38y). The scale in grossly comparable with that of the LCA4 patients that did not have axial length measurements. Arrows highlight the trace ellipsoid zone in the retina of the LCA4 patients. OCT revealed residual foveal outer retinal structure in all three children. All three children had limited visual acuity to perception of light. (NFL = nerve fibre layer, GCL = ganglion cell layer, IPL = inner plexiform layer, INL = inner nuclear layer, OPL = outer plexiform layer, HFL = Henle’s fiber layer, ONL = outer nuclear layer, ELM = external limiting membrane, EZ = ellipsoid zone, IZ = interdigitation zone, RPE = retinal pigment epithelium).

**Supplementary Figure 2:** Reprogramming of LCA4 patient renal epithelial cells and characterisation of iPSC lines.

A) Brightfield images of LCA4 patient renal epithelial cells which were reprogrammed to generate iPSCs. Emerging iPSC colonies were isolated and iPSC lines established from single clones. Scale bars = 250μm.

B) IF analysis of LCA4-1, LCA-3 and LCA-4 iPSCs for the expression of pluripotency markers (OCT4, Nanog, SSEA4, TRA-1-60, TRA-1-81). DAPI staining is in blue. Scale bars = 100μm.

C) Trilineage analysis of iPSC lines. Real time PCR (qPCR) analysis of iPSC lines subjected to ectodermal (*PAX6*, *NESTIN*, *OTX2*)/ mesodermal (*T*)/ endodermal (*CDX2, AFP, SOX17, GATA6*) culture conditions. The pluripotency marker *OCT4* was downregulated in differentiation cultures. All expression levels normalised to housekeeping β-actin gene. Values = mean +/− SD.

**Supplementary Figure 3:** Additional characterisation of LCA4 patient ROs.

A) Brightfield images of developing retinal organoids (D40-230) from LCA4-2, LCA4-3 and LCA4-4 patient iPSC lines. Scale bars = 250μm.

B) TUNEL staining of W28 CTL and LCA4 RO sections. Very few fluorescein (green) positive cells were observed, and these were located in the core of the ROs, where non-PR retinal cell types reside. DAPI staining is in blue. Scale bars = 100μm.

C) IF of W28 RO sections for AIPL1 (polyclonal antibody raised against recombinant full-length AIPL1). This alternate AIPL1 antibody confirmed that LCA4 ROs lack detectable levels of AIPL1 protein. DAPI staining is in blue. ONL and INL regions highlighted at the side of images in blue and magenta. Scale bars = 50μm.

**Supplementary Figure 4:** Characterisation of c.834G>A, p.W278X CRISPR-Cas9 HDR-repaired LCA4-1 isogenic iPSC lines.

A) Brightfield images of LCA4 isogenic (isogenic lines 1 (iso.1) and 2 (iso.2)) iPSC colonies. Scale bars = 250μm.

B) IF of LCA4-1 iPSC colonies (iso.1 and iso.2) for pluripotency markers (OCT4, TRA-1-60, TRA-1-81). DAPI staining is in blue. Scale bars = 100μm.

C) Trilineage assay analysis of LCA4 isogenic iPSCs (iso.1 and iso.2). qPCR results showed upregulation of germ-layer specific markers and downregulation of pluripotency marker, *OCT4*. All expression levels normalised to housekeeping β-actin gene. Values = mean +/− SD.

**Supplementary Figure 5:** Analysis of AIPL1 translational readthrough levels in LCA4-1 ROs dosed with varying amounts of PTC124 and effect on retinal cell type lineages.

LCA4-1 ROs were treated with 5-15ug/ml PTC124 for 2 weeks, prior to IF analysis. Treatment with 10-12.5ug/ml of PTC124 appeared to be the most efficient at increasing the levels of AIPL1 protein in LCA4-1 ROs. DAPI staining is in blue. ONL and INL regions are highlighted at the side of images in blue and magenta. Scale bars = 50μm.

**Supplementary Figure 6:** Further analysis of LCA4-1 ROs: IF and TUNEL assay.

A) IF of W24 LCA4-1 isogenic, LCA4-1 untreated and LCA4-1 PTC124-treated ROs for the expression of AIPL1, Rhodopsin (RHO), Transducin (GT1, rod), and cone opsin (OPN1SW). With the exception of AIPL1, the PR markers were similarly distributed in all ROs. DAPI staining is in blue. ONL and INL regions are highlighted at the side of images in blue and magenta. Scale bars = 50μm.

B) TUNEL staining of W24/28/33 RO sections showed that apoptosis rates are similar in all RO types at all timepoints with low levels of TUNEL positive cells. DAPI staining is in blue. ONL and INL regions are highlighted at the side of images in blue and magenta. Scale bars = 50μm.

C) Semi-quantitative gene expression analyses of LCA4-1 isogenic control, LCA4-1 untreated and PTC124-treated ROs at different developmental timepoints (W12-W33) for genes relating to retinal development and retinal cell lineages.

D) Quantitative PCR analysis of *CRX* levels in LCA4-1 isogenic, LCA4-1 untreated and 10ug/ml PTC124-treated ROs (W12-W33, 3 biological replicates per timepoint). No significant differences were found between sample types within timepoints. Gene expression levels normalised to housekeeping gene, β-actin (*ACTB*).

## Tables

Supplementary Table 1:

List of Antibodies

Supplementary Table 2:

List of Primers – AIPL1 Genotyping / Sanger sequencing

Supplementary Table 3:

List of Primers for gene expression analyses – qPCR / semi-quantitative PCR

Supplementary Table 4:

List of sgRNAs and ssODN Sequences – CRISPR/Cas9 HDR

Supplementary Table 5:

List of Primers – CRISPR/Cas9 HDR Predicted Off-Target sites.

