## Supplementary Data for "Translational readthrough as a potential therapeutic for AIPL1-associated Leber Congenital Amaurosis in a patient-derived iPSC-retinal organoid model"

### Supplementary Figure 1

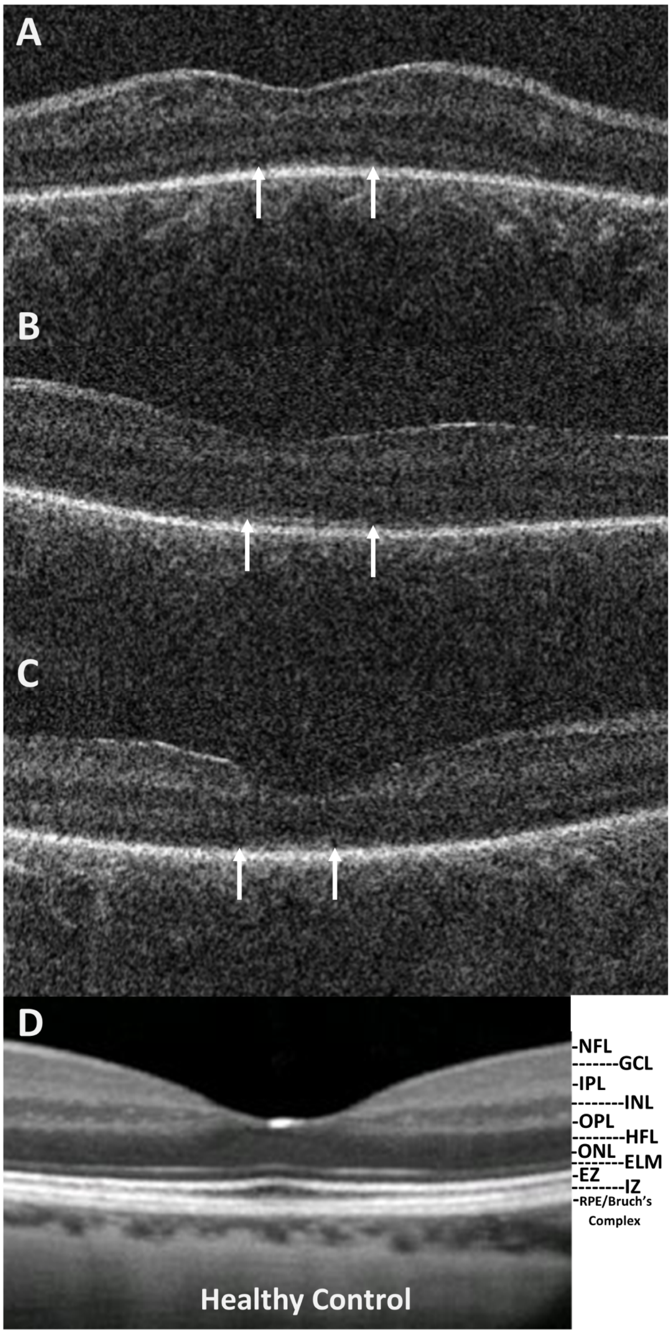

### Supplementary Figure 2

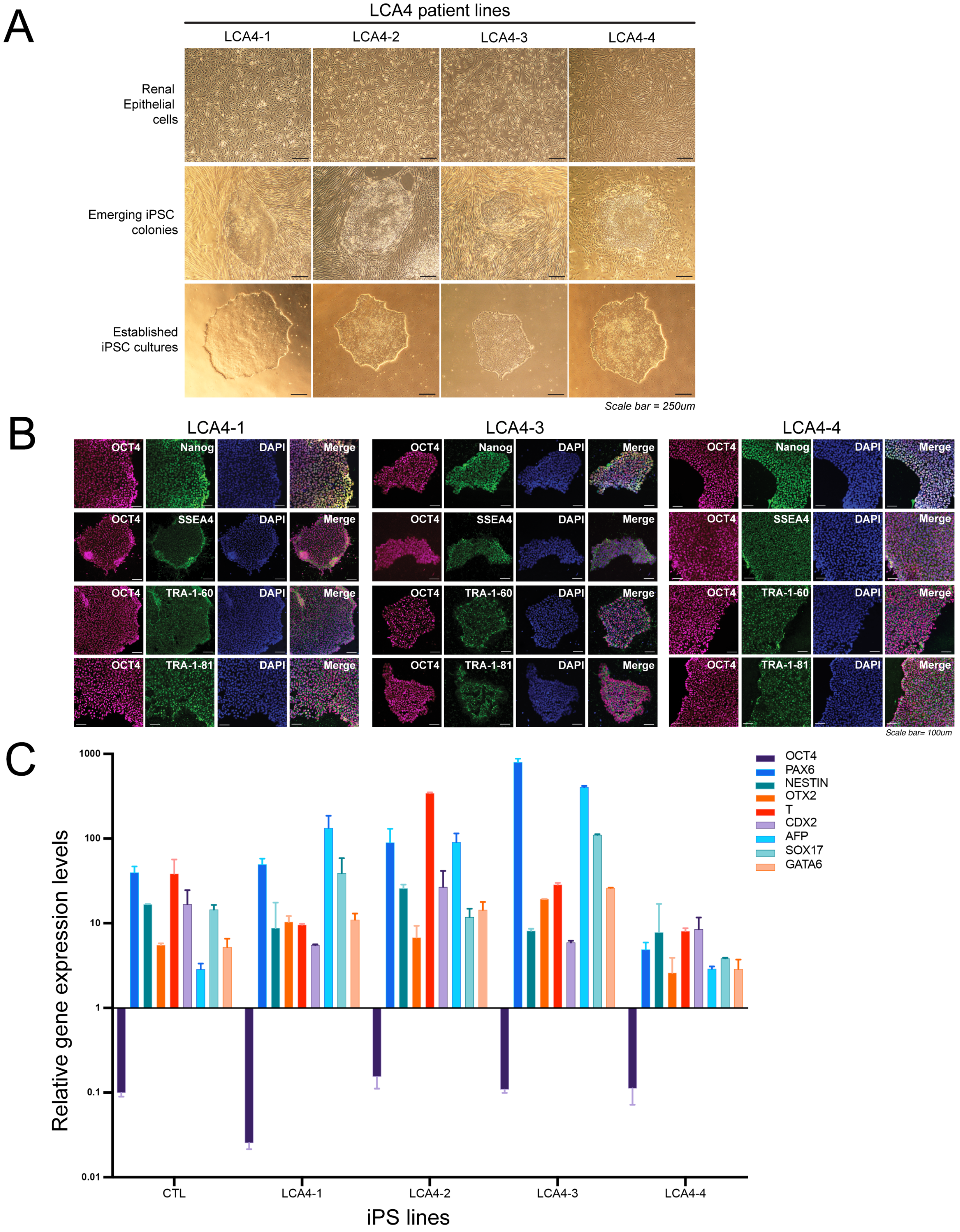

### Supplementary Figure 3

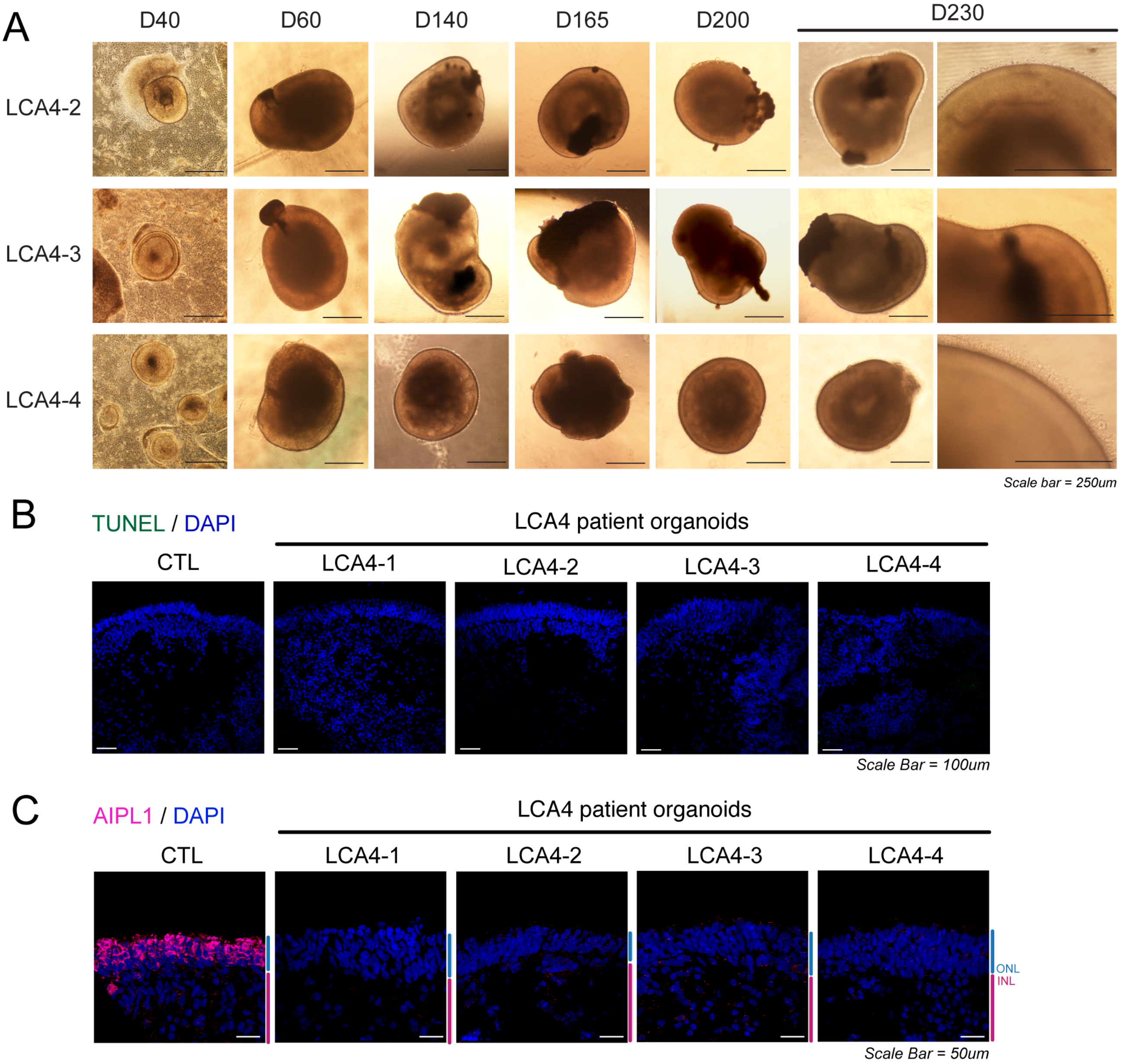

### Supplementary Figure 4

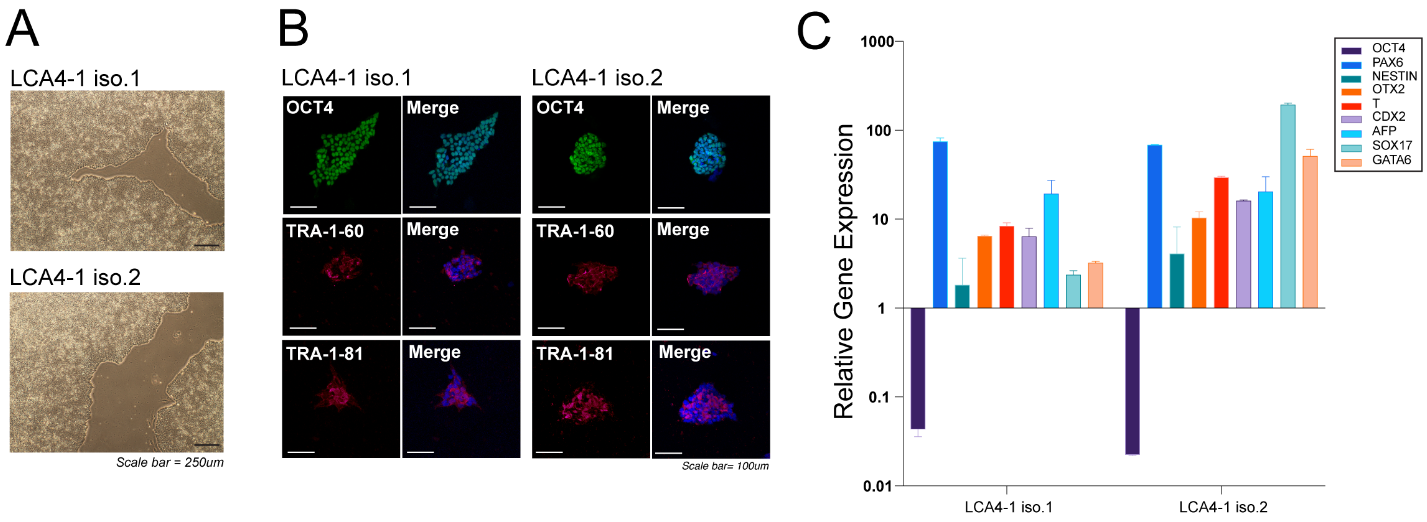

Supplementary Figure 5

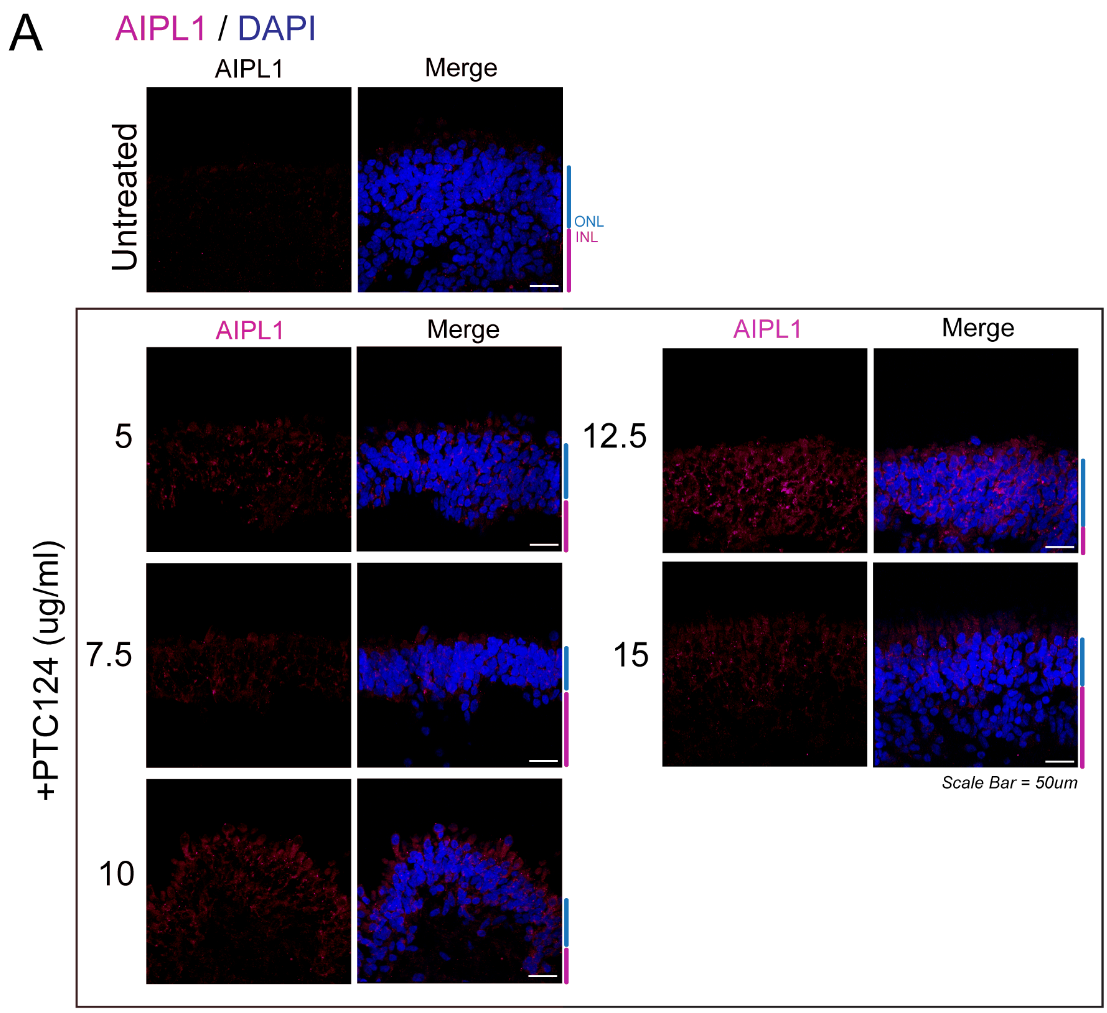

### Supplementary Figure 6

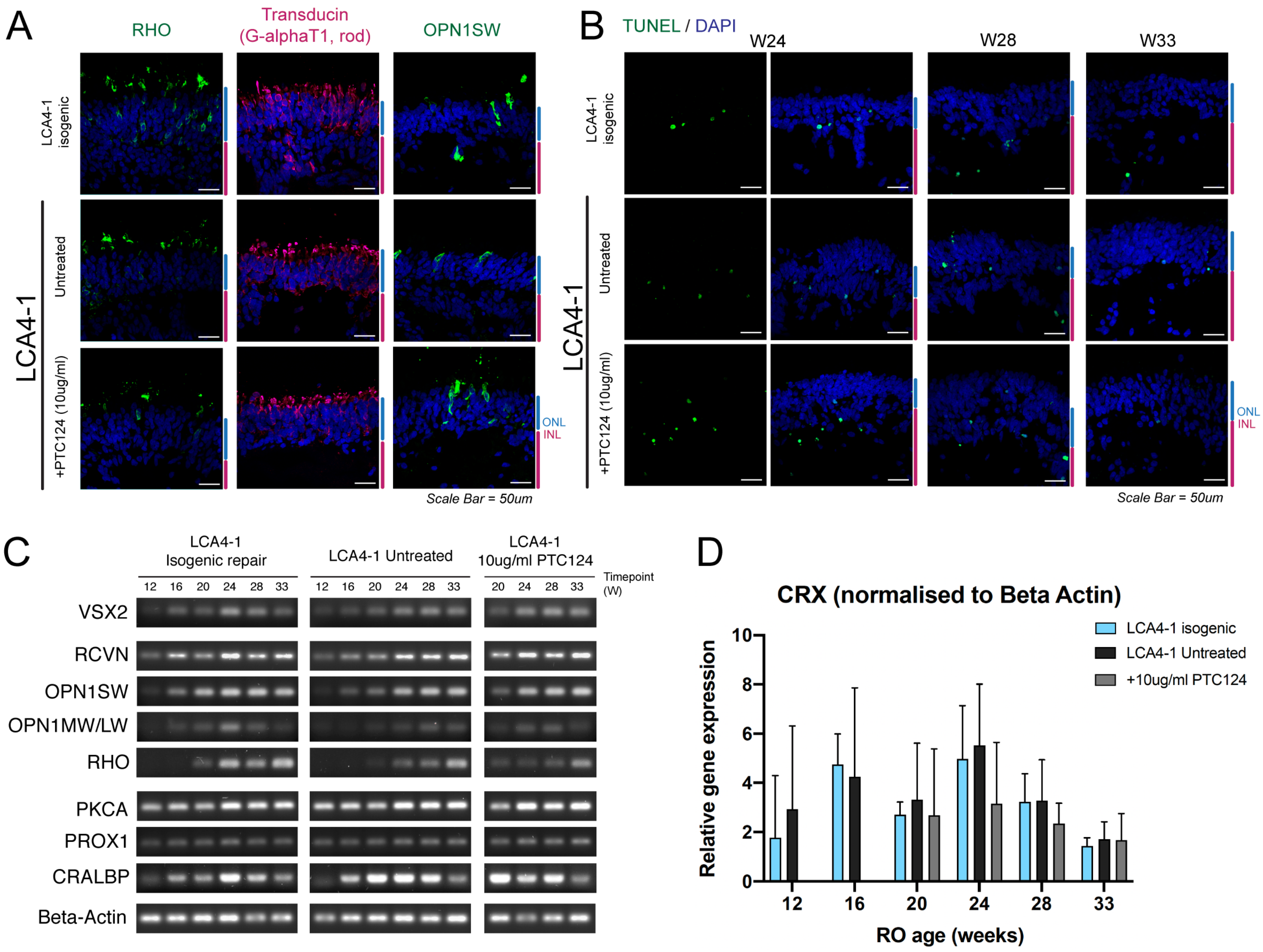

### Supplementary Table 1:

**ICC : antibodies and stains used**

| **Antibody** | **Species** | **Antibody info** | **Catalogue no. / AB** | **Dilution** |
|---|---|---|---|---|
| AIPL1 | Rabbit | J. van der Spuy lab, raised against AIPL1 C-ter | | 1 / 250 |
| AIPL1 | Rabbit | V. Ramamurthy lab | | 1 / 500 |
| PDE6A | Rabbit | Abcam | ab5659 | 1 / 1000 |
| PDE6B | Rabbit | Thermo Fisher Scientific | PA1-722 | 1 / 1500 |
| cGMP | Sheep | BioRad | OBT5055 | 1 / 100 |
| RETGC1 | Rabbit | A.M. Dizhoor lab | / | 1 / 4000 |
| Rhodopsin | Mouse | Millipore | 4D2 | 1 / 1000 |
| Cone Arrestin | Mouse | 7G6 monoclonal AB | / | 1 / 200 |
| RTLN | Mouse | Santa Cruz | C-20 / sc-374056 | 1 / 200 |
| L/M-Opsin | Rabbit | Millipore | AB5405 | 1 / 500 |
| S-Opsin | Rabbit | Millipore | AB5407 | 1 / 500 |
| Transducin, Rod (Gat1) | Rabbit | Santa Cruz | K-20 / sc-389 | 1 / 100 |
| Recoverin | Rabbit | Millipore | | 1 / 500 |
| PKCA | Rabbit | Abcam | ab32376 | 1 / 2000 |
| ISLET1 | Rabbit | Santa Cruz - discontinued | / | 1 / 50 |
| BRN3B | Rabbit | Santa Cruz - discontinued | / | 1 / 10 |
| Vimentin | Rat | R and D systems | MAB2105-SP | 1 / 500 |
| PROX1 | Rabbit | Sigma Aldrich | AB5475 | 1 / 2000 |
| OCT4 | Rabbit | Abcam | ab19857 | 1 / 1000 |
| NANOG | Rabbit | Abcam | ab21624 | 1 / 1000 |
| SSEA4 | Mouse | Cell signaling | MC813 / #4755 | 1 / 1000 |
| TRA-1-81 | Mouse | Cell signaling | #4745 | 1 / 1000 |
| TRA-1-60 | Mouse | Cell signaling | #4746 | 1 / 1000 |

| **Secondary Antibodies** | **Fluorophore** | **Company** | **Catalogue no. / AB** | **Dilution** |
|---|---|---|---|---|
| Donkey anti-mouse IgG | AF488 | Thermo Fisher Scientific | A-21202 | 1 / 1000 |
| Donkey anti-mouse IgG | AF555 | Thermo Fisher Scientific | A-31570 | 1 / 1000 |
| Donkey anti-rabbit IgG | AF555 | Thermo Fisher Scientific | A-31572 | 1 / 1000 |
| Donkey anti-rabbit IgG | AF488 | Thermo Fisher Scientific | A-21206 | 1 / 1000 |
| Donkey anti-sheep IgG | AF488 | Molecular Probes | / | 1 / 1000 |
| Donkey anti-rat IgG | AF488 | Thermo Fisher Scientific | A-21208 | 1 / 1000 |

| **Stain** | **Fluorophore** | **Company** | **Catalogue no. / AB** | **Dilution** |
|---|---|---|---|---|
| Phalloidin | AF488 | Invitrogen / Molecular Probes | A12379 | 1 : 40 of 200U/ml solution |

### Supplementary Table 2

**AIPL1 genotyping/sequencing primers**

| Amplicon | Region | Forward primer | Reverse primer | Expected Size |
|---|---|---|---|---|
| Genomic | **Exon 1** | ACTGGAAGCAAAGGTGGAT | CCATGCTAAAGTTGAATCTG | 526bp |
| Genomic | **Exon 2** | TGAACTGAGTGAGCTGACCC | GAATAAGTTTGCAGGACTGGCTTTG | 428bp |
| Genomic | **Exon 3** | CATAGTGAGGGAGCAGGATTC | CATGGCTTATGAACCCTCTCG | 441bp |
| Genomic | **Exon 4** | CTTGTCTGTATGCACTTGACCAG | CAGGGAGAAGGTCAGCCATG | 426bp |
| Genomic | **Exon 5** | CGGCTGGGTGGAGACAAG | GAAGTGGCGCTGACTCTGG | 369bp |
| Genomic | **Exon 6** | TTGAGGAAACCGAGGGATGG | CAATCGAACCAGAAGTGACCAGG | 582bp |
| Transcript | **Ex3-5** | TCTACCCCATCCTATCCCG | GGAGAATATCACTGGTGTGCTC | 486bp |
| Transcript | **Ex5-6** | CTGATCCTCAACTACTGCCAG | AGGTGGCTCTGTGGATGA | 360bp |

Supplementary Table 3

**Real Time PCR and Semi-quantitative PCR**

| Marker | Forward primer | Reverse primer | Notes |
|---|---|---|---|
| **AIPL1** | ACCGGATCCCGAGTGATCTT | CGATGATGATGTGCATGGGC | qPCR/semi-quant |
| **VSX2** | GTGGCTACTGGGGATGCAC | TCCTGCTCCATCTTGTCGAG | Semi-quant |
| **PAX6** | AACGATAACATACCAAGCGTGTC | GTCTGCCCGTTCAACATCCT | Semi-quant |
| **NRL** | CACTGACCACATCCTCTCGG | GAGGGTTCCCGCTTTACCTC | Semi-quant |
| **CRX** | TTTGCCAAGACCCAGTACC | GTTCTTGAACCAAACCTGAACC | qPCR/semi-quant |
| **Recoverin** | ATGAAGTGCTGGAGATCGTC | ATCTTCTCGGCTCGCTTTTC | Semi-quant |
| **Rhodopsin** | ACCAGCACCCTCTACACCTCTC | AGGACCACAGGGCAATTTC | Semi-quant |
| **OPN1SW** | CATGTTTGTGCTTTGGAGG | CGAAGGGCTTACAGATGAC | Semi-quant |
| **MW/LW-OPN1** | CCTATGTGTGTCCTGGAGG | CATCCATCTCTCCCAGGAAATG | Semi-quant |
| **RETGC1** | ACTGTCCCTCTGAAGGCAG | CGTCATAGATGGTGCCAAAG | qPCR/semi-quant |
| **PDE6A** | TAACGTCCCCAACACAGAGG | CCACCACATCCTTCCCATTC | qPCR/semi-quant |
| **PDE6B** | GACGTGTGGTCTGTGCTGAT | CTTGCCGTGGAGGATGTAGTC | qPCR/semi-quant |
| **PDE6G** | AAGCAGCGACAGACCAGG | TGTGATGTCTGTTCCCAGGC | Semi-quant |
| **PDE6C** | GTCACTAAGAACCTGCTGGCAACC | AAAGACCTCTTCATCCTGTTTGG | Semi-quant |
| **PDE6H** | GAGGCAGACTCGCCAATTC | GTGGCTGAATGCCTCCCA | Semi-quant |
| **PKCA** | GTCCACAAGAGGTGCCATGAA | AAGGTGGGGCTTCCGTAAGT | Harvard Primer Bank / Semi-quant |
| **ISL1** | GCGGAGTGTAATCAGTATTTGGA | GCATTTGATCCCGTACAACCT | Harvard Primer Bank / Semi-quant |
| **CRALBP** | AAGCTGGCTACCCTGGTGT | TGAAGCAATATGCCTGCAAGA | Harvard Primer Bank / Semi-quant |
| **BRN3B** | CTCGCTCGAAGCCTACTTTG | GACGCGCACCACGTTTTTC | Harvard Primer Bank / Semi-quant |
| **PROX1** | TGAAGACCTACTTCTCCGAC | GACGTGCGTACTTCTCCATC | Semi-quant |
| **OCT4** | TTTGCCAAGCTCCTGAAGCA | AAGGGCCGCAGCTTACACAT | qPCR |
| **PAX6** | GCGGTGAGAAGTGTGGGAAC | GCCCGTTGACAAAGACACCA | qPCR |
| **Nestin** | TCAGATGTGGGAGCTCAATCG | GCTCTTCAGCCAGGTTGTCG | qPCR |
| **OTX2** | CGCAGTCAATGGGCTGAGTC | ACCGGGTCTTGGCAAACAGT | qPCR |
| **Brachyury (T)** | CCTTCAGCAAAGTCAAGCTCACC | TGAACTGGGTCTCAGGGAAGCA | qPCR |
| **CDX2** | TCCTGGACAAGGACGTGAGC | CGCGTAGCCATTCCAGTCCT | qPCR |
| **AFP** | TGAGCACTGTTGCAGAGGAG | TTGTTTGACAGAGTGTCTTGTTGA | qPCR |
| **SOX17** | GGATACGCCAGTGACGACCA | CTCGTCCTTAGCCCACACCA | qPCR |
| **GATA6** | CTGAACGGGACGTACCACCA | GTCTGGATGGAGCCGCAGTT | qPCR |
| **GAPDH** | CCCCACCACACTGAATCTCC | GGTACTTTATTGATGGTACATGACAAG | qPCR/semi-quant |
| **Beta Actin** | CCAACCGCGAGAAGATGA | CCAGAGGCGTACAGGGATAG | qPCR/semi-quant |

### Supplementary Table 4

#### List of sgRNAs and ssODN Sequences – CRISPR/Cas9 HDR

| Combination | sgRNA sequence | ssODN template sequence* (* denote phosphorothioate (PS) bonds) |
|---|---|---|
| 1 | TCACGCAGAGGTGTGAAATG | T*C*CAGCAGCCTCAGCTCCCTGCGCACCGCCTTCTGCATGGACGGCTCCAGCTCCAGCACTTTCTGGAGGTCCGCCTT GGCCTCGGCTTCGTTCCACACCTCTGCGTGAGCCCGGGCACGCACGTA*G*T |
| 2 | AGAGGTGTGAAATGAGGCCG | C*G*GTTCTCCAGCAGCCTCAGCTCCCTGCGCACCGCCTTCTGCATGGACGGCTCCAGCTCCAGCACTTTCTGGAGGTC CGCCTTGGCCTCGGCCTCGTTCCACACCTCAGCGTGAGCCCGGGCACG*C*A |

The combination of sgRNA1 and ssoDN template1 was able to trigger CRISPR-Cas9 HDR of the p.W278X locus with an estimated efficiency of editing at the p.W278X locus of approximately 30%, with HDR enhancer (as determined by TIDER analysis (http://shinyapps.datacurators.nl/tider/)). This was used to establish LCA4-1 isogenic iPS lines.

No editing was detected with sgRNA2 and ssODN template2.

### Supplementary Table 5

#### AIPL1-W278X locus CRISPR: off-target analysis

| Program | Region | Site of potential off-target editing | Sequence of potential binding site* | Bulge size* | Mismatches* | Forward primer | Reverse primer |
|---|---|---|---|---|---|---|---|
| CASOFFFINDER | **Site 1** | Chr.14, NC_000014.9: 67687177 to 67687198 | crRNA: TCACGCAGAGGTGTGAAATGNGG<br>DNA: TCA-GCAGAGGTGaGAAATGAGG | 1 | 1 | CACCTAAAGCATGCATCTCC | CTAAGAGCAATGGGAAACGG |
| CASOFFFINDER | **Site 2** | Chr.8, NC_000008.11 : 142474973 to 142474996 | crRNA: TCAC-GCAGAGGTGTGAAATGNGG<br>DNA: TCACAGCtGgGGTGTGAAATGTGG | 1 | 2 | AGGCCTTGATCACCACTG | CAGGCTCCGTCCAGTTTC |
| CASOFFFINDER | **Site 3** | Chr.3, NC_000003.12 : 172174110 to 172174131 | crRNA: TCACGCAGAGGTGTGAAATGNGG<br>DNA: gCAtGCAGAGGTGTGAAA-GTGG | 1 | 2 | GCGAACTGAGAATCCATTCATG | CCAAATATGCTTGAATGCCTTG |
| CASOFFFINDER | **Site 4** | Chr.4, NC_000004.12 : 186616865 to 186616886 | crRNA: TCACGCAGAGGTGTGAAATGNGG<br>DNA: TaA-GCAGAGGTGTtAAATGGGG | 1 | 2 | GTTGGATGCAACTGGAGTC | TCAATGGGACCATCATTTGG |
| CASOFFFINDER | **Site 5** | Chr.7, NC_000007.14 : 146187291 to 146187314 | crRNA: TCAC-GCAGAGGTGTGAAATGNGG<br>DNA: TCACAGCAGAGGTGTtAcATGTGG | 1 | 2 | CCAACTCTGGTAAATGTGAGTAG | CTCTTCTTCGGCTCTCAGTG |
| OFFSPOTTER | **Site 6** | Chr.10, NC_000010.11 : 131402426 to 131402448 | crRNA: TCACGCAGAGGTGTGAAATGNGG<br>DNA: gCggGCAGAGGTGTGAAATGAGG | 0 | 3 | TCTTAATATTGATGCCAGCCTG | AGCCAGTTCTCCTGTTGAT |
| OFFSPOTTER | **Site 7** | Chr.X, NC_000023.1: 25154339 to 25154361 | crRNA: TCACGCAGAGGTGTGAAATGNGG<br>DNA: aCAaGCAGAGaTGTGAAATGTGG | 0 | 3 | AATGTACTTCACTACACATCGTC | GTCAACTCAACATTTCTCTCTTC |
| OFFSPOTTER | **Site 8** | Chr.4, NC_000004.12 : 186616865 to 186616887 | crRNA: TCACGCAGAGGTGTGAAATGNGG<br>DNA: TtAaGCAGAGGTGTtAAATGGGG | 0 | 3 | Same as for Site 4 | Same as for Site 4 |
| OFFSPOTTER | **Site 9** | Chr.4, NC_000004.12 : 21759877 to 21759899 | crRNA: TCACGCAGAGGTGTGAAATGNGG<br>DNA: TCAaGCAGAGGTGaaAAATGAGG | 0 | 3 | CCACCTGTCCTTGTAAACTG | GTTGGAGACCTGTAGTTCTTTC |
| OFFSPOTTER | **Site 10** | Chr.12, NC_000012.12 : 59076201 to 59076223 | crRNA: TCACGCAGAGGTGTGAAATGNGG<br>DNA: gaAaGCAGAGaTGTGAAATGCGG | 0 | 4 | GCTGAAATAATCATGAAGGCATG | CAGATATAGCTGAATCCAGGTAC |

***Bulges and base pair mismatches are highlighted in red**
